# Lysophosphatidylcholine induces heat pain hypersensitivity in obese mice fed with a high-fat diet through activation of peripheral Acid-Sensing Ion Channel 3

**DOI:** 10.1101/2021.12.07.471593

**Authors:** Negm Ahmed, Stobbe Katharina, Fleuriot Lucile, Debayle Delphine, Deval Emmanuel, Lingueglia Eric, Rovere Carole, Noel Jacques

## Abstract

Diet induced obesity is one of the major causes of obesity, which affects 13% of the world’s adult population. Obesity is correlated to chronic pain regardless of other components of the metabolic syndrome. Our study focuses on investigating the effect of high-fat diet induced obesity on peripheral sensory neurons activity and pain perception, followed by deciphering the underlying cellular and molecular mechanisms that involve Acid-Sensing Ion Channel 3 (ASIC3). We show here that heat sensitive C-fibers from mice made obese by consumption of a high-fat diet exhibited an increased activity during baseline and upon heating. Obese mice showed long-lasting heat pain hypersensitivity once obesity was well established, while mechanical sensitivity was not affected. We found that the serum of obese mice was enriched in lysophosphatidylcholine (LPC) species (LPC16:0, LPC18:0 and LPC18:1), which activate ASIC3 channels and increased peripheral neuron excitability. Genetic deletion and *in vivo* pharmacological inhibition of ASIC3 protected and rescued mice from obesity-induced thermal hypersensitivity. Our results identify ASIC3 channels in DRG neurons and circulating LPC species that activate them as a mechanism contributing to heat pain hypersensitivity associated with high-fat diet induced obesity.

## Introduction

The prevalence of obesity is a growing public health problem worldwide affecting 13% of the adult population. Obesity and overweight, measured as high body mass index (BMI > 30 and 25, respectively) are well established risk factors for chronic diseases, such as cardiovascular disease, diabetes mellitus, kidney diseases, cancer and chronic musculoskeletal pain (Afshin et al., 2017). The main cause of obesity is the imbalance between the energy intake provided by food and the energy expenditure through physical activity. This leads to the accumulation of excessive body fat and a chronic metabolic inflammatory state or low-grade chronic inflammation in peripheral tissues, which is responsible for other comorbidities. Chronic inflammation also reaches the nervous system and the brain, particularly the hypothalamic nuclei that control feeding behavior (Gregor & Hotamisligil, 2011). Growing evidence suggests that deleterious alterations in lipid metabolism contribute to this chronic inflammatory state. Specifically, altered serum levels of pro-inflammatory lipids, such as arachidonic acid and lysophosphatidylcholine (LPC), have been related to obesity (Barber et al., 2012; Graessler et al., 2009; Pickens et al., 2017; Pietiläinen et al., 2007).

Pain is a common comorbidity of obesity (Mills et al., 2019). It is one of the most pervasive symptom and one of the leading causes of disability and disease burden worldwide (Vos et al., 2017). Studies on human subjects suggest that obesity is an independent predictor of pain at all ages (Hainsworth et al., 2009; Hitt et al., 2007; Marcus, 2004; Mills et al., 2019; Webb et al., 2003). Obese and overweight people are more likely than others to experience all types of pain, which further supports a link between obesity and pain (Hitt et al., 2007; Okifuji & Hare, 2015; Ray et al., 2011). Preclinical studies using animal models have also shown a strong association between obesity and pain (Rodgers et al., 2014; Song et al., 2017). However, the causative relationship between both disorders is not clear.

A growing body of evidence suggests that pro-inflammatory and anti-inflammatory lipid mediators play a crucial role in the hypersensitization of nociceptors associated with inflammation. We have shown that lysophosphatidylcholine (LPC) and arachidonic acid (AA) are endogenous activators of Acid-Sensing Ion Channel 3 (ASIC3) at resting physiological pH (Deval et al., 2008; Marra et al., 2016). The combination of both lipids activate nociceptive C-fibers in the skin and induced acute pain behaviors in rodents that were dependent on ASIC3 (Marra et al., 2016). More recently, elevated levels of LPC detected in the synovial fluids of patients with painful rheumatic diseases were correlated with pain outcomes, and this lipid was found to trigger ASIC3-dependent chronic joint pain and anxiety-like behaviors when injected intra-articularly in mice (Jacquot et al., 2021). ASIC3 channels (Waldmann et al., 1997), which belong to the family of degenerin/epithelial sodium channels (DEG/ENaC), are predominantly expressed in neurons of sensory ganglia, including DRG (Lin et al., 2016; Wu et al., 2012; Yan et al., 2013). They contribute to nociceptor sensitization through the activation of a voltage-independent non-inactivating current upon extracellular acidification (Deval et al., 2003) or exposure to lipid activators (Deval et al., 2008). They have been involved in processes related to pain (Deval & Lingueglia, 2015) such as acid-induced pain in the skin and muscle (Deval et al., 2008; Walder et al., 2010), post-operative pain (Deval et al., 2011), fibromyalgia (Hsu et al., 2019; Hung et al., 2020), and in acute and sub-acute inflammatory pain (Deval et al., 2008; Karczewski et al., 2010; Yen et al., 2009).

We were therefore interested in investigating whether elevated lipid serum concentration may contribute to the sensitization of nociceptors in the context of obesity, and the possible contribution of ASIC3 in this effect. We show that high-fat diet (HFD)-induced obesity in wild-type mice is associated with a state of pre-diabetes and induces thermal hypersensitivity related to ASIC3 channel activity. Application of serum collected from obese HFD-fed mice increased the excitability of cultured DRG neurons and skin nociceptors sensitivity to heat. Lipidomic analysis of the serum from obese HFD-fed mice showed elevated levels of the pro-inflammatory lipid lysophosphatidylcholine (LPC). Delipidation of the serum from HFD-fed mice reversed its activity on DRG neurons, while supplementation of delipidized serum with LPC restored its effect. Genetic deletion or pharmacological inhibition of ASIC3 prevented the excitation of DRG neurons and corrected HFD-induced thermal hyperalgesia. DRG neurons from ASIC3-null mice were not activated by application of serum from HFD-fed obese mice. ASIC3-null mice on HFD became obese but did not show thermal hyperalgesia.

## Materials and Methods

All animal procedures were approved by the Institutional Local Ethical Committee and authorized by the French Ministry of Research (Agreements 02677.01) according to the European Union regulations and the Directive 2010/63/EU and was in agreement with the guidelines of the Committee for Research and Ethical Issues of the International Association for the Study of Pain (Zimmermann, 1983). Animals were sacrificed at experimental end points by CO_2_ euthanasia.

### Animals and diet

Experiments were performed on wild-type (WT) male C57Black6J mice (Janvier labs, France) and ASIC3 knockout mice (ASIC3 ko) (Wultsch et al., 2008). Four weeks-old mice 15-18 g were used for the feeding experiments. Mice were weight-matched between groups and acclimated to housing and husbandry conditions for at least one week before experiments. Animals were housed in 12 hours reversed light-dark cycle (lights on at 20:00). One group received control standard diet (SD) and the other received high-fat diet (HFD), *ad libitum* access to food and water. The weight of the mice, the amount of food and water consumed were recorded 3 times a week.

The two diets, SD and HFD, are from Scientific Animal Food and Engineering (SAFE®). The SD is a standard breeding diet given to the control mice. The caloric intake of the SD is 2.830 kcal/kg, mainly through proteins and fibers (25.4% of total nutritional composition) and with only 5.1% from lipids. Lipids in SD correspond to an equilibrated balance between polyunsaturated (linoleic acid, omega-6, and linolenic acid, omega-3), mono-unsaturated (oleic acid) and saturated fatty acid (palmitic acid) (Supplementary Table 1). The ratio ω-6/ ω-3 is 6.3. The high-fat diet with high concentration of saturated lipids from butter was used to produce obese mice. HFD contains high concentration of lipids, 34-36% in diet, which is designed to increase total body weight and adipose body mass, with diabetogenic effect. HFD caloric intake is 5.283 kcal/kg. HFD brings 61 % of the food energy from fats, mainly from saturated fatty acids (69 % of fat). Palmitic and stearic acids account for 45% of fat. The ratio ω-6/ω-3 is 5.6. Less than 1% of fat content is from ω-3 poly-unsaturated fatty acids (alpha-linolenic acid).

### Glucose (GTT) and Insulin (ITT) Tolerance tests

Mice were respectively fasted 16h (GTT) and 6h (ITT) prior to intraperitoneal (IP) injection of glucose (1.5 g/kg, Sigma-Aldrich) or insulin (0.75 U/kg, Novo Nordisk)(Ayala et al., 2010). Glycemia was then measured by tail blood sampling at 0, 15, 30, 60, 90 and 120 min post injection using a glucometer (Accuchek Performa, Roche). During the experiments, mice were unrestrained and were freely moving.

### Nociceptive behavior assays

For the nociceptive behavior experiments (Deuis et al., 2017), mice were placed in transparent observation chambers where they were acclimated for at least 20 min.

### Radiant heat Hargreaves test

Thermal hyperalgesia was measured with radiant heat Hargreaves test (Bioseb, France). Mice were placed in transparent plastic boxes on an elevated floor. A radiant heat source was then placed under one animal’s hind paw and maintained until the mouse lifted its paw (cutoff time 20 s). Each measure was repeated twice on both hind paws, with more than 3 min interval between measures.

### Tail immersion test

Heat sensitivity was tested by immersing the tail of the mice in a temperature-controlled water bath at 46°C until tail withdrawal was observed (cutoff time 30 s). Mice were habituated for two days, and measurements were done on the third day. The latency of tail withdrawal was calculated by averaging 3 responses separated by at least 3 min.

### Thermal perception assessment with pharmacological inhibition of ASIC3

Two groups of 8 WT mice received either SD or HFD. Thermal perception was evaluated with Hargreaves tests performed at 4 and 7 weeks of diet. Following the evaluation at 7 weeks, mice from both groups (SD- and HFD-fed mice) received a total of 7 IP injections of the peptide APETx2 (0.23 μg/g), an ASIC3 inhibitor (Diochot et al., 2004), every 2 days over 2 weeks. For injections, mice were gently restrained while APETx2 was administered IP using a 26-gauge needle connected to a 100-ml Hamilton syringe. Hargreaves test were performed over the period of APETx2 treatment, at 8 weeks of diet (*i.e.*, after 4 APETx2 injections) and at 9 weeks of diet (*i.e.*, after 7 injections). Tests were then performed after stopping the treatment, at 12 weeks (*i.e.*, 3 weeks after the last APETx2 injection) and at 16 weeks of diet (*i.e.*, after 7 weeks from the last injection).

In paralleled experiments, 2 groups of 8 WT mice received either SD or HFD. Thermal perception was evaluated with Hargreaves tests at 4, 8, and 12 weeks of diet before APETx2 injections in both groups. APETx2 (0.23μg/kg) was injected IP every 2 days over 2 weeks. Hargreaves test were performed over the injection period at 7, 9, 12, 14 days post injections.

Mechanical sensitivity. The mechanical sensitivity was evaluated with a dynamic plantar aesthesiometer test (Bioseb, France). Mice were placed in individual plastic boxes on top of a wire surface. A filament was applied with increasing force, ramped up to 7.5 g in 10 sec, on the plantar surface of mice hind paws until a paw withdrawal response was elicited. The paw withdrawal force threshold (g) was measured in duplicate for each paw at more than 3 min interval. Mechanical sensitivity was also assessed using von Frey filaments ranging from 0.02 to 1.4g (Bioseb, France). Mice were placed in individual plastic boxes on a wire mesh surface, von Frey filaments of increasing stiffness were applied 4 times, at more than one-minute interval, to the plantar surfaces of the hind paw. The mechanical threshold was evaluated as the strength of filament that elicited more than 2 responses.

### Chemicals

APETx2 was synthesized by Synprosis/Provepep (France). Lipids were purchase from Sigma and Avanti (Coger, France). All lipids were prepared as stock solutions in EtOH, stored at −20°C, and diluted to the final concentration extemporaneously before the experiments. The final concentration of EtOH was below 0.12 %.

### Serum preparation and delipidation

After blood collection from mice, serum was obtained by centrifugation at 10,000g, for 5 min at 4°C in Micro tube 1.1ml Z-Gel (Sarstedt, Germany). Samples were kept on ice at all times. The protocol for serum delipidation was adapted from (Renaud et al., 1982) where the serum density was adjusted with NaBr to be 1.215 g/ml using Radding-Steinberg formula. Serum was then ultra-centrifuged at 223,000g for 24 h and the lower phase containing the delipidated serum was collected. This step was repeated 3 times. The delipidized serum was dialyzed in Slide-A-Lyzer® Dialysis cassette (Thermo scientific), cutoff of 3.5 kDa, against control bath solution containing (in mM): 145 NaCl, 5 KCl, 2 MgCl_2_, 2 CaCl_2_, and 10 HEPES (pH 7.4 with NaOH). This procedure reduced the concentration of lipids present in HFD serum to levels close to the concentration measured in serum from SD-fed mice (Fig. 4A). Of note, the depletion was homogeneous among LPC species. Serum pH was monitored before application at 7.69 +/− 0.03 for HFD serum and 7.62 +/− 0.06 for SD serum (n=13-5 mice respectively, p= 0.2, unpaired t-test). Osmolarity was monitored at 360 +/− 10 mOsm for HFD-S and 341 +/− 3 mOsm for SD-S (n=9-5 mice, p=0.19, unpaired t-test).

### Lipidomic mass spectrometric analysis

#### Lipids extraction

Lipids were extracted according to a modified protocol from (Bligh & Dyer, 1959). Lipid extraction was performed in 1.5 ml solvent-resistent plastic Eppendorf tubes and 5 ml glass hemolyse tubes to avoid contamination. Methanol, chloroform, and water were kept at 4°C. 120 μL of serum was collected in a 1.5 ml Eppendorf tube and 300 μl of methanol was added. After vortexing (30s), the sample was frozen for 20 min at −20°C. The sample was transferred in a glass tube and 100 μL of chloroform was added. The mixture was vortexed for 30s and centrifuged (2,500 rpm, at 4°C for 10 min). After centrifugation, the supernatant was collected and 150 μL of chloroform and 150 μL of water were added. The sample was vortexed again for 30s and then centrifuged (2,500 rpm, 4°C, and 10 min). 150 μL of the non-polar phase was collected and dried under a stream of nitrogen. The dried extract was re-suspended in 60 μL of methanol/chloroform 1:1 (v/v) and transferred in an injection vial before liquid chromatography and mass spectrometry analysis.

**Reverse phase liquid chromatography** was done with an Ultra Performance Liquid Chromatography (UPLC) system (Ultimate 3000, ThermoFisher). Lipid extracts of serum from mice were separated on an Accucore C18 150×2.1, 2.5μm column (ThermoFisher) operated at 400 μl/min flow rate. The injection volume was 3 μl of diluted lipid extract. Eluent solutions were Acetonitrile (ACN)/H_2_O 50/50 (V/V) containing 10mM ammonium formate and 0.1% formic acid (solvent A) and isopropropanol/ACN/H_2_O 88/10/2 (V/V) containing 2mM ammonium formate and 0.02% formic acid (solvent B). Acetonitrile and isopropropanol were purchased from Carlo Erba, chloroform from Merck, methanol from VWR. Formic acid Optima LC-MS quality was purchased from ThermoFisher, ultrapure water from purelab flex (Veolia Water), and formate ammonium (99%) from Acros Organics. The step gradient was: 0.0–4.0 min 35% to 60% B, 4.0–8.0 min 60 to 70% B, 8.0–16.0 min 70 to 85% B, 16.0-25 min 85 to 97% B, 25-25.1 min 97 to 100% B, 25.1-31 min 100% B and finally the column was reconditioned at 35% B for 4.0 min. The UPLC system was coupled with a Q-exactive orbitrap Mass Spectrometer (ThermoFisher, CA) equipped with a heated electrospray ionization (HESI) probe. The spectrometer was controlled by the Xcalibur^TM^ software and operated in electrospray positive mode. MS spectra were acquired at a resolution of 70 000 (200 m/z) in a mass range of 250−1.200 m/z with an AGC target 1e6 value and a maximum injection time of 250 ms. The 15 most intense precursor ions were selected and isolated with a window of 1 m/z and fragmented by HCD (Higher energy C-Trap Dissociation) with normalized collision energy (NCE) of 25 and 30 eV. MS/MS spectra were acquired in the ion trap with an AGC target 1e5 value, in a mass range of 200-2000 m/z, the resolution was set at 35 000 m/z combined with an injection time of 80 ms. Data were reprocessed using Lipid Search 4.1.16 (Thermo Fisher). The identification was based on the accurate mass of precursor ions and MS2 spectral pattern. Mass tolerance for precursor and fragments was set to 5 ppm and 8 ppm respectively. M-score threshold was selected at 5 and ID quality filter was fixed at grades A, B and C. [M+H]+, [M+Na]+ and [M+NH4]+ adducts were searched.

**LPC quantification** was done using a standard calibration curve. For that, we used three LPC standards: LPC16:0, LPC18:0 and LPC18:1. Each standard was analyzed with six different concentrations (5, 10, 25, 40, 50, and 70 μM). Each concentration was injected 4 times for technical replicates with the same UPLC-HRMS method used for serum analysis. The measures given in arbitrary unit (area under the peak) were fitted against their corresponding standard concentrations by linear regression.

### Cultures of DRG neurons

DRG neurons from WT mice were prepared from male C57Bl6J mice or ASIC3 knockout mice (8 to 14 weeks old) based on previous study (Noël et al., 2009). After isoflurane anesthesia, animals were killed by decapitation and lumbar DRG were collected on cold HBSS and enzymatically digested at 37°C for 40 min with collagenase II (Biochrom, 2mg/ml, 235U/mg). Gentle mechanical dissociation was done with a 1ml syringe and several needles with progressive decreasing diameter tips (18G, 21G and 26G) to obtain cell suspension. Neurons were then plated on collagen-coated 35 mm Petri dishes (Biocoat) and maintained in culture at 37°C (95% air/5% CO_2_) with the following medium: Neurobasal-A (Gibco) completed with L-glutamine (Lonza, 2mM final), B27 supplement and 1% peniciline/streptomycine (Gibco). One day after plating, neurons were carefully washed to remove cellular debris and incubated with complete medium including additional growth factors: Nerve Growth Factor (NGF) 100ng/ml, retinoic acid 100nM (Sigma), Glial Derived Neurotrophic Factor (GDNF) 2ng/ml, Brain Derived Neurotrophic Factor (BDNF) 10ng/ml and Neurotrophin 3 (NT3) 10ng/ml (all these from Peprotech). Patch-clamp recordings were done 1-4 days after plating on small to medium diameter neurons, having a cell membrane capacitance <40 pF, the majority of which are considered to be nociceptive neurons (Bardoni et al., 2014).

### HEK-293 cell cultures and transfection

HEK-293 cell line was grown as described previously (Marra et al., 2016). One day after plating, cells were transfected with 0.5μg of DNA per dish of one of the following plasmids, pIRES2-mASIC3-EGFP or pIRES2-rASIC1a-EGFP vectors using the JetPEI reagent according to the supplier’s protocol (Polyplus transfection SA, Illkirch, France). Fluorescent cells were used for patch-clamp recordings 2–4 days after transfection.

### Patch-clamp experiments

The whole-cell configuration of the patch-clamp technique was used to measure membrane currents and voltages (voltage-clamp and current-clamp configuration respectively). Recordings were performed at room temperature using an axopatch 200B amplifier (Axon Instruments) with a 2 kHz low-pass filter. Recordings were sampled at 10 kHz, digitized with a Digidata 1440 A-D/D-A converter (Axon Instruments) and recorded on a hard disk using pClamp software (version 10; Axon Instruments). The patch pipettes (2–6 MΩ) contained (in mM): 135 KCl, 2.5 Na_2_-ATP, 2 MgCl_2_, 2.1 CaCl_2_, 5 EGTA, and 10 HEPES (pH 7.25 adjusted with KOH). The control bath solution contained (in mM): 145 NaCl, 5 KCl, 2 MgCl_2_, 2 CaCl_2_, 10 HEPES, and 10 glucose for neurons, pH adjusted with N-methyl-D-glucamine or NaOH. ASIC currents were induced by shifting one out of eight outlets of a micro-perfusion system from a control solution (*i.e.*, pH 7.4) to acidic test solution at pH 6.6 or 7.0.

To measure the effect of the serum on DRG neuron’s rheobase, DRG neuron membrane potentials were maintained at −55 mV with patch-clamp current-clamp configuration and steps of increasing current were injected in the pipette until triggering the first action potentials in control conditions and after perfusing the DRG neurons with serum collected from either the SD or the HFD mice. The fold change in rheobase was calculated by dividing the current required to trigger the rheobase during the serum application with the rheobase during the control period. The threshold of action potential was determined as the voltage that corresponds to the first derivative of the action potential at 10 mV/ms. The capacitance and resting membrane potential of DRG neurons used within each group are listed in supplementary Tables 2 and 4.

### Nerve-skin preparation and single fiber recordings

Single C-fiber recording technique from the isolated skin-saphenous nerve preparation was used as previously described (Pereira et al., 2014). After euthanasia the hind paw skin of mouse (8-12 weeks on diet; 12 – 16 week-old mice) was dissected with the saphenous nerve. The skin was superfused with warm (32°C) synthetic interstitial fluid (SIF) (in mM): 120 NaCl, 3.48 KCl, 5 NaHCO_3_, 1.67 NaH_2_PO_4_, 2 CaCl_2_, 0.69 MgSO_4_, 9.64 Na-gluconate, 5.5 glucose, 7.6 sucrose, and 10 HEPES, pH adjusted to 7.4 with NaOH, saturated with O_2_/CO_2_ - 95%/5%. The receptive field of an identified C-fiber was searched by mechanical probing of the skin and further characterized for its conduction velocity (below 1.3 m/s) and mechano-sensitivity with calibrated von Frey filaments. A fiber was classified as a C-fiber when its conduction velocity was less than 1.3 m/s. This protocol implies that all C-fibers were mechano-sensitive. C-fibers’ receptive fields were isolated with a thick walled stainless steel elrin ring (internal volume of 400 μl) inside which solutions were applied through local perfusion pipes of a CL-100 bipolar temperature controller (Warner instrument). The perfusion solution was kept at 32°C for baseline and raised to 48-50°C in 50 sec during a heat-ramp. Recordings were band-pass filtered between 60 Hz and 2 kHz and sampled at 10 kHz on computer with pClamp 10 software (Axon Instrument). The analysis of action potentials was done with Spike2 software (Cambridge Electronic Design) and spikes were discriminated off-line and visualized individually.

### Data statistical analysis

Data analysis was performed using Microcal Origin 8.5, R studio and GraphPad Prism 4.03 softwares. Data are presented as mean ± s.e.m. Statistical differences between sets of data were assessed using either parametric or nonparametric tests followed by post-hoc tests, when appropriate. D’Agostino and Pearson normality tests were done for all groups to check normality. Unpaired t-test or Mann-Whitney test were used for unpaired comparisons, while Wilcoxon matched-pairs signed rank test was used for paired comparisons. Kruskal-Wallis test followed by Dunn’s multiple comparisons test was used for multiple group comparisons and 2-way ANOVA with repeated measures followed by Bonferroni’s multiple comparisons test was used whenever appropriate.

## Results

### High fat diet led to obesity and a state of pre-diabetes until 16 weeks of diet

To investigate the comorbidity between nutritional obesity and pain hypersensitivity, we fed four weeks-old juvenile C57black6/J wild-type male mice (WT) with a high-fat diet (HFD; 34% of fat from butter) that provides 60% of dietary energy from lipids with significantly higher caloric intake per week (Fig 1A). Within four weeks of regime, HFD-fed mice gained significantly more weight than mice fed with a standard diet (SD), which brings only 5% of dietary energy from lipids (Fig 1B). Weight difference between HFD- and SD-fed mice increased during the whole period of regime, such that after 16 weeks of diet, the weight of HFD-fed mice was on average 37% higher than SD-fed mice (37.6 +/− 1.9 g and 27.5 +/− 1 g respectively; p< 0.0001 2-way ANOVA with repeated measures followed by Bonferroni’s post hoc test, n=10 per group).

**Fig. 1.**
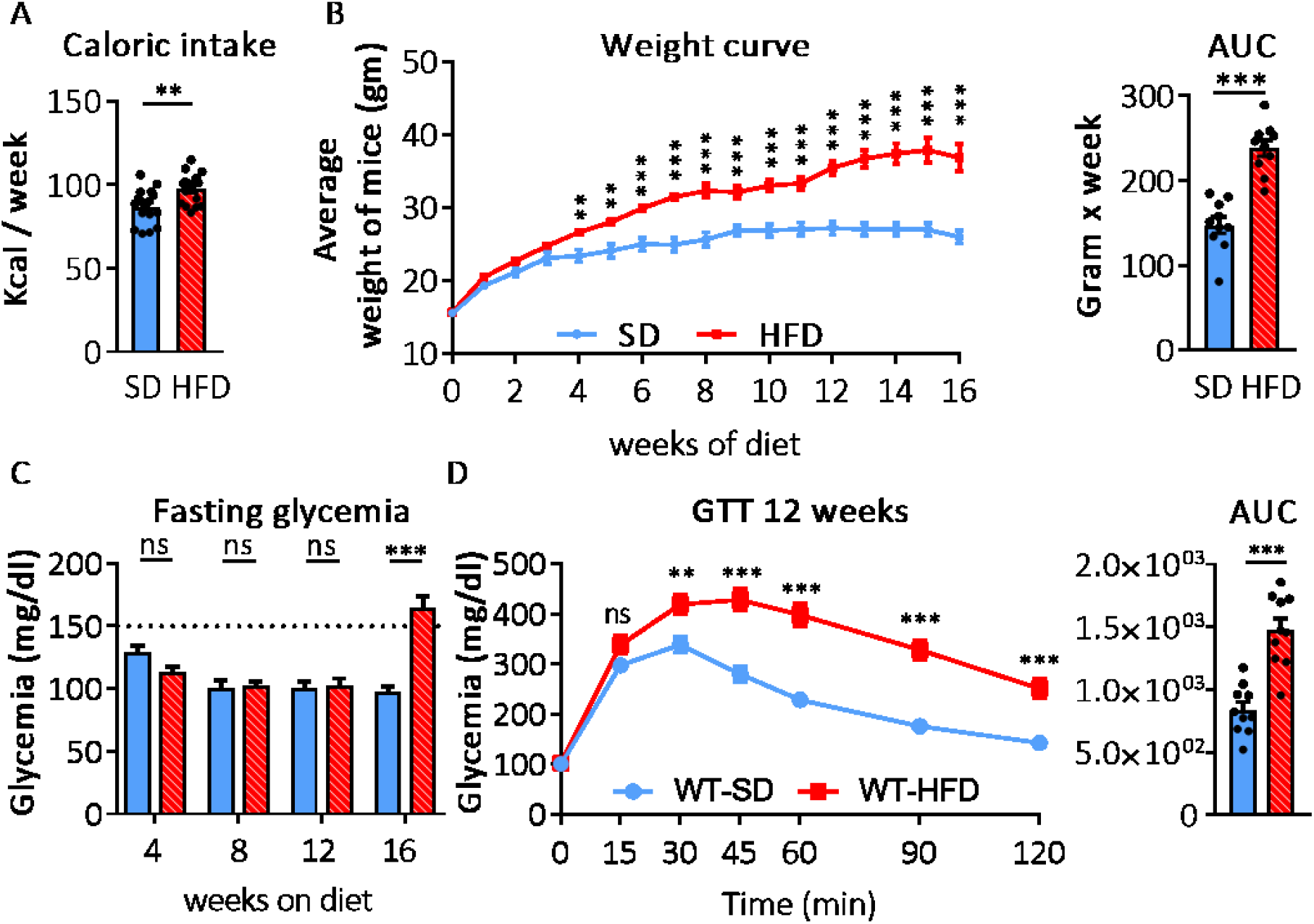
Lipid-rich diet consumption induces obesity and a prediabetes state. **(A)** Caloric intake of wild-type (WT) mice fed high-fat diet in (red) and WT mice fed the standard diet (SD) (blue) (n=10 per group, **p= 0.005; unpaired t-test). Color codes are conserved all over the figure. **(B)** Weight gain over weeks on diet for WT-SD and WT-HFD mice (n=10 per group, F (16, 288) = 10.48, p< 0.0001, asterisks indicate p-values; 2-way ANOVA with repeated measures followed by Bonferroni’s post-hoc test). Right panel: area under the curve (AUC) of the total weight gain during the whole feeding period (16 weeks) (n=10 per group, ***p<0.0001, unpaired t-test). **(C)** Fasting glycemia measured at different weeks of regime as are indicated for the SD-fed and the HFD-fed mice (n= 10 per group, ***p< 0.0001, 2-way ANOVA with repeated measures followed by Bonferroni’s post-hoc test). **(D)** Glucose tolerance test (GTT) at 12 weeks of diet for WT mice fed with either SD or HFD (glycemia at 120 min post injection was 251.4 +/− 19.3 mg/dl and 143.3 +/− 4.9 mg/dl for HFD- and SD-fed mice respectively, p<0.0001, n=10 per group; asterisks indicate P-values obtained from 2-way ANOVA with repeated measures followed by Bonferroni’s post-hoc test). Right panel is the AUC (*** p<0.001, unpaired t-test). Data are presented as mean ± SEM. ns = not significant; *, ** and *** for p values <0.05, <0.01 and <0.001 respectively.

We monitored glucose metabolism over a period of 16 weeks of diet and we observed that it was progressively deregulated by HFD. Fasting blood glucose concentrations remained below 125 mg/dl during the first 12 weeks of regime and were not different between HFD- and SD-fed WT mice (Fig. 1C). After 16 weeks of diet, however, HFD-fed mice showed hyperglycemia with a fasting glucose concentration significantly higher than the one of SD-fed mice (164.3 +/− 9.7 mg/dl and 97.8 +/− 4 mg/dl for HFD-fed and SD-fed mice respectively; p < 0.0001, repeated measures 2-way ANOVA followed by Bonferroni’s post hoc test, n=10 per group). The fasting blood glucose concentration of HFD-fed mice after 16 weeks of diet was above 150 mg/dl, which could indicate an early phase of diabetes (O’Brien et al., 2014; Obrosova et al., 2007). The glucose tolerance test (GTT) showed a progressive deregulation of glucose metabolism in HFD-fed WT mice compared to SD-fed mice. After 4 weeks of HFD, mice started to show a slight alteration in glucose metabolism (Supplementary Fig. 1A). However, after 8 and 12 weeks of diet both the maximal blood glucose concentration and the concentration reached 2 hours after a bolus injection of glucose were increased in HFD-fed compared to SD-fed mice (Fig1D and Supplementary Fig. 1B). At 16 weeks of diet, the dysregulation in glucose homeostasis evolved and the difference in glycemia between mice fed with the HFD and the SD was significantly different at all-time points (Supplementary Fig. 1C). In addition, HFD-fed mice also developed an increased resistance to insulin within the first 12 weeks of diet (Supplementary Fig. 5D). Deregulation of glucose homeostasis and increased insulin resistance are indicators of Type 2 diabetes during the period before 16 weeks of diet, and we focused our studies on mice that were on diets for 8 to 12 weeks showing signs of an early phase of diabetes mellitus or pre-diabetes.

### Mice fed the lipid-rich diet developed thermal hypersensitivity and increased heat-sensitive C-fibers activity

We then examined whether HFD-induced obesity was associated with pain hypersensitivity in mice. Mechanical pain tested with dynamic von Frey and von Frey filaments showed no difference between HFD- and SD-fed WT mice when tested after 8, 12 or 16 weeks of regime (Fig. 2A, Supplementary Fig.2). Conversely, HFD-fed mice showed a hypersensitivity to heat when tested with Hargreaves or tail-flick tests (Fig. 2B, C). A significant reduction of PWL was measured with the Hargreaves test in HFD-fed mice after only 8 weeks of regime (at 8 weeks of diet, p= 0.0001, 2-way ANOVA with repeated measures followed by Bonferroni’s post-hoc test, n= 10 per group; Fig. 2B). This was confirmed in the tail-flick test at 46°C with reduction of tail flick latencies for mice on HFD for 8 weeks (Fig.2C). Heat-pain hypersensitivity of HFD-fed mice was preserved over 20 weeks of diet, while over the same period the heat response of SD-fed mice did not fluctuate.

**Fig. 2.**
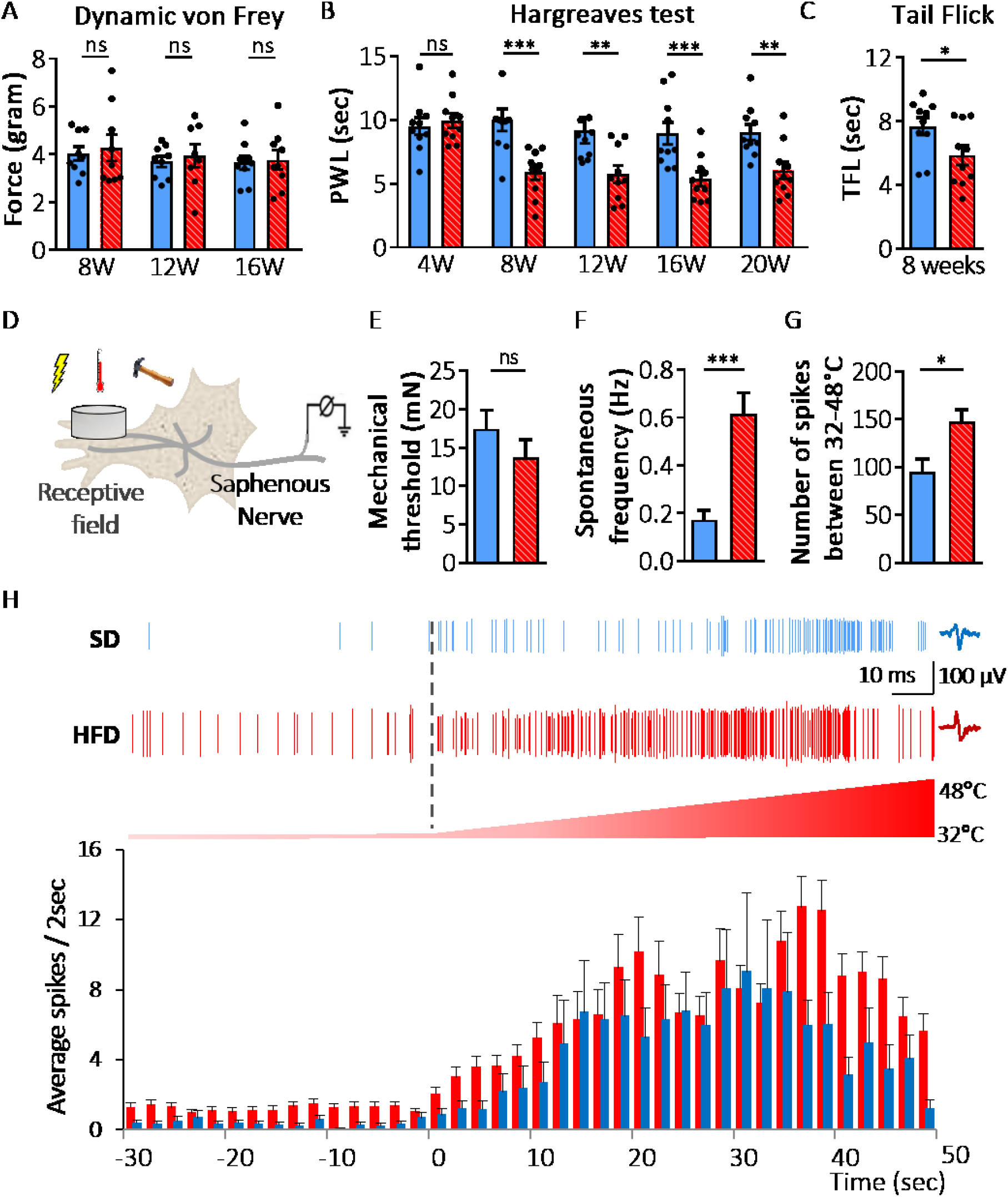
HFD induces heat-thermal hyperalgesia without affecting mechanical perception. **(A)** Mechanical perception of obese HFD-fed mice compared to control SD-fed mice presented as grams using the dynamic von Frey test at different weeks of diets (F (2, 31) = 0.02, p=0.98, 2-way ANOVA with repeated measures followed by Bonferroni’s post-hoc test n= 9 per group). Red bars for HFD-fed and blue bars for SD-fed mice. **(B)** Paw withdrawal latencies measured with Hargreaves test for SD-fed and HFD-fed WT mice over time (n=10 per group, ns= not significant, p>0.05, ** p<0.01, *** p<0.001, 2-way ANOVA with repeated measures followed by Bonferroni’s post hoc tests). **(C)** Tail flick experiment at 46°C at 8 weeks of diet on lean SD-fed and obese HFD-fed WT mice. Tail flick latencies were 7.7 +/− 0.6 sec and 5.9 +/− 0.6 sec for SD-fed and HFD-fed mice respectively (n=10 per groups, p=0.04, unpaired t-test). **(D)** Cartoon representing the *ex-vivo* skin-saphenous nerve preparation with elrin ring placed on the recorded fiber’s receptive field in the skin. **(E)** Mean mechanical threshold of mechanosensitive C-fibers (mN) from SD- and HFD-fed mice (average mechanical thresholds were 17.5 +/− 2.4 mN and 13.8 +/− 2.3 mN, respectively; n=12-15 C-fibers from SD and HFD mice recorded from 4-6 animals; p=0.2, Mann Whitney test). **(F)** Mean spontaneous firing frequency (Hz) of heat sensitive C-fibers at 32°C recorded during 30 s from SD- and HFD-fed mice (mean spontaneous frequency were 0.2 +/− 0.04Hz and 0.6+/− 0.08Hz for C-fibers from SD and HFD mice respectively; n=23-48 C-fibers from 4-6, ***p= 0.0007, Mann Whitney test). **(G)** Average total number of action potentials of heat sensitive C-fibers during a heat ramp from 33°C to 48°C in 50 sec (averaged total number of spikes were 95 +/− 13.5, and 148 +/− 12.8; n= 23-48 C-fibers from 4-6 SD and HFD mice respectively, p=0.02 Mann Whitney test). **(H)** Top panel: representative firing activity of exemplar single C- fibers from SD- and HFD-fed mice during a heat ramp. Normal temperature of the skin (32°C) and during warming of the skin to 48°C. The grey dotted line indicates the beginning of the heat ramp. The temperature ramp is presented below the 2 traces with a range from 32°C to 48°C. Lower panel shows the cumulative activity of heat sensitive C-fibers of skin from SD- and HFD-fed mice presented as the mean number of spikes during 2 seconds plotted against time, corresponding to 30 s baseline at 32°C followed by a heat ramp reaching 48°C in 50 sec (n= 23-48 C-fibers from 4-6 SD- and HFD-fed mice respectively).

We then aimed to investigate the neuron-based mechanisms that were responsible for the heat hypersensitivity of HFD-fed obese mice. Heat pain is initiated by the stimulation of nociceptive C-fibers innervating the skin, and therefore, we measured the activity of mechanical and heat-sensitive cutaneous C-fibers of HFD-fed and SD-fed mice, on diet for 8 to 12 weeks, with the *ex-vivo* nerve-skin recording technique (Zimmermann et al., 2009)(Fig 2D-H). The receptive fields of individual C-fibers in the hind-paw skin of WT mice were probed for mechanical sensitivity with calibrated von Frey filaments while action potentials activity was recorded. We measured no difference of mechanical sensitivity between C-fibers of HFD- and SD-fed mice (Fig.2E). We next measured the response of individual C-fibers to heat by increasing the temperature of the skin that enclose their receptive field from the physiological temperature of 32°C to a painful temperature of 48°C within 50 s. First, we found that the spontaneous activity of C-fibers from HFD-fed mice was higher than SD-fed mice at the physiological temperature of the skin of 32°C (Fig. 2F). Furthermore, when the skin temperature was increased to 48°C, C-fibers of HFD-fed mice fired more action potentials than C-fibers from SD-fed mice (Fig. 2G). The elevated activity was observed over the entire range of temperatures (Fig. 2H, lower panel). These nerve-skin recordings showed that C-fibers of HFD-fed mice were hypersensitive to heat but not to mechanical stimuli. The response of nociceptive C-fibers from the skin was consistent with pain hypersensitivity to heat and not to mechanical stimuli that was observed *in vivo* in HFD-fed obese mice. These results suggest that peripheral mechanisms that affect heat-sensitive nociceptive C-fibers are involved in the thermal sensitization induced by HFD consumption, although not excluding possible central mechanisms.

### Serum from obese mice excites peripheral sensory neurons through activation of ASIC3 channels

We next sought to investigate the causal link between HFD-induced obesity and sensory neuron hyper-excitability. Peripheral neurons innervating organs like skin are not protected by the blood brain barrier (Feldman et al., 2017), and therefore, changes in the composition of serum after the consumption of HFD could directly alter sensory neurons activity. To test this hypothesis, we first decided to measure *in vitro* the activity of small to medium diameter dorsal root ganglion neurons (DRGs) in cultures prepared from WT mice, while perfusing them with serum collected from obese HFD-fed mice (HFD-S). The activity of the neurons was recorded in the current-clamp mode of the patch-clamp whole-cell configuration (Fig. 3A). Interestingly, the application of HFD-S depolarized the membrane potential of all DRG neurons tested with mean depolarization of 7.9+/-1.1 mV (Fig.3A, right panel). Moreover, the application of HFD-S triggered the firing of action potentials in 3 out of 9 recorded neurons (Fig. 3A, left panel) highlighting that some components of the HFD serum excite DRG neurons.

**Fig. 3.**
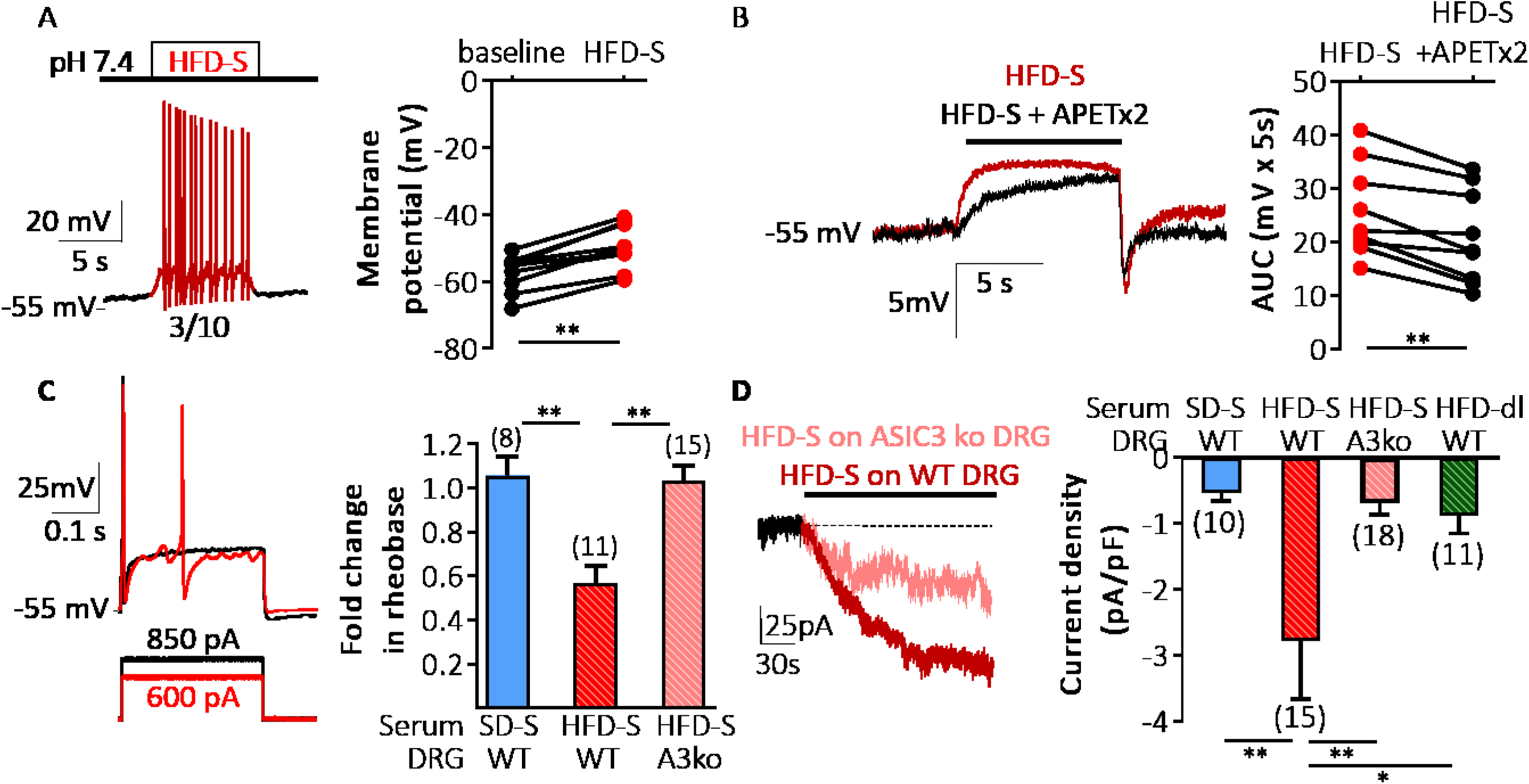
HFD serum increases excitability of DRG neurons through ASIC3 channels. **(A)** Left panel shows a representative recording, in patch-clamp current-clamp configuration, of a DRG neuron from WT mouse perfused with HFD serum (HFD-S). Right panel shows the depolarizing effect of HFD-S (red dots) on WT-DRG neurons compared to their resting membrane potential (black dots) (mean membrane potential were −54.7+/−2.5 mV and −49.5+/−2.2 mV for pre and during the application of the HFD serum, n=9 neurons, p=0.004, Wilcoxon matched-pairs signed rank test). Mean capacitance was 48.9+/−5.3pF. **(B)** Left panel shows two superimposed representative current-clamp recordings of a WT DRG neuron during the application of HFD-S alone (red trace) or HFD-S with the ASIC3 inhibitor APETx2 (2μM; HFD-S+APETx2) applied for 1 min prior and during the application of the serum (black trace). Right panel shows the depolarization induced by both solutions calculated as the area under the curve measured during 5 sec of application (mean depolarization over 5 sec were 25.7 +/− 2.9 mV and 20.8 +/− 2.9 mV for HFD-S and HFD-S with APETx2, n=9, p=0.004, Wilcoxon test. Capacitance and membrane resting potentials as in Fig. 3A. **(C)** Effect of serum on DRG neurons rheobase. Left panel shows two superimposed representative recordings at the rheobase of the membrane potential of a DRG neuron without serum (black trace) and during the application of HFD-S (red trace). Right panel shows the fold change in the rheobase obtained upon application of SD-S and HFD-S on WT DRG neurons (1.06 +/− 0.09 and 0.56 +/− 0.08 respectively) and application of HFD-S on DRG neurons from ASIC3 knockout mice (HFD A3 ko, 1.03 +/− 0.07, number of neurons recorded are indicated above bars; cell membrane capacitances and resting membrane potentials are listed in Supplementary table 2; p= 0.009 HFD-S vs SD-S both applied on WT-DRGs, p= 0.001 HFD-S applied on WT-DRGs vs ASIC3 ko DRGs; Kruskal-Wallis test followed by Dunn’s multiple comparisons test). **(D)** Left panel shows two superimposed representative whole-cell currents, recorded at −80 mV in patch-clamp voltage-clamp configuration, of the effect of HFD-S application on DRG neurons cultured from WT (red trace) and ASIC3 knockout mice (pale red trace). Right panel shows the mean calculated current densities of currents evoked by the application of SD-S, HFD-S and delipidized HFD-S (HFD-dl) on WT DRGs (current densities: −0.5 +/− 0.1 pA/pF, −2.8 +/− 0.9 pA/pF and −0.9 +/− 0.3 pA/pF, respectively), and HFD-S on ASIC3 knockout DRG neurons (HFD-S A3 ko, current density: −0.7 +/− 0.2 pA/pF, number of neurons recorded are indicated above each bar; current densities are measured 60 sec after serum application; values of resting membrane potentials and cell membrane capacitances are listed in Supplementary Table 4; p= 0.006 HFD-S vs SD-S both applied on WT DRGs, p= 0.002 HFD-S applied on WT DRGs vs ASIC3 ko DRGs, p=0.04 for HFD-S vs HFD-dl both applied on WT DRGs, p>0.9 SD-S vs HFD-dl both applied on WT DRGs; Kruskal-Wallis test followed by Dunn’s multiple comparisons test).

ASIC3 channels expressed in DRG neurons produce a depolarization of the membrane upon extracellular acidification or the application of lipids, notably AA and LPC (Deval et al., 2008; Poirot et al., 2006). This raised the interesting possibility that ASIC3 channels expressed in DRG neurons could support the depolarizing effect of HFD-S. To test this hypothesis, we used APETx2, a peptide toxin from sea anemone that blocks ASIC3 channels (Diochot et al., 2004), to investigate the contribution of ASIC3 to this effect. We measured that the depolarization induced by HFD-S was significantly reduced by the co-application of 2 μM APETx2 (Fig. 3B, right panel). This observation supports a role for ASIC3 channels in the excitation of DRG neurons by serum from HFD-fed mice, although an additional contribution of Nav channels to the effect, which were shown to be inhibited to some extent by APETx2 (Peigneur et al., 2012), cannot be fully excluded.

To further investigate how HFD-S affected DRG neurons excitability, we measured its effect on the rheobase required to trigger action potentials. For that, we used the patch-clamp recording in the whole-cell current-clamp configuration to inject current steps of increasing intensity in DRG neurons and we compared the current amplitude that triggered the first action potential during the application of serums collected from SD- or HFD-fed mice (Fig. 3C). Membrane potential was set at −55 mV to counter the depolarization triggered by HFD serum and to be able to compare measurements from different neurons. The rheobase of WT DRG neurons was not changed by the application of serum from SD-fed mice (SD-S) (Fig. 3C, right panel). On the contrary, the application of serum collected from HFD-fed mice significantly lowered the rheobase of WT DRG neurons (Fig. 3C). Interestingly, the threshold for action potential triggered by the rheobase current was not changed by the application of SD or HFD serum (Supplementary Table 3). To confirm the possible contribution of ASIC3 channels in neuronal hyperexcitability induced by HFD-S, we compared the rheobase of DRG neurons from ASIC3 knock-out mice (ASIC3 ko) with and without the application of HFD-S (Fig. 3C). The serum collected from HFD mice did not change the rheobase of DRG neurons cultured from ASIC3 ko mice nor the action potential threshold (Fig. 3C and Supplementary Table 3). These experiments show that the serum of HFD-fed mice activates and induces hyperexcitability of small to medium diameter DRG neurons and that ASIC3 is one of the main mediators of this effect. Several ion channels other than ASIC3 might be involved in the neuronal excitability induced by HFD-S including Na^+^ and K^+^ channels. Therefore, to minimize possible contribution of these channels we measured the effect of serum on whole-cell currents of cultured DRG neurons in the whole-cell voltage-clamp recording configuration at a holding potential of −80 mV. This holding potential is the reversal potential for K^+^ and will exclude the contribution of K^+^ channels, and it is well below the activation thresholds of voltage-dependent Na^+^ channels, thus favoring the investigation of the voltage-insensitive Na^+^ permeable channels like ASIC3. The application of HFD-S on WT DRG neurons activated a sustained inward current that was active as long as the serum was present (Fig. 3D). The current evoked by HFD-S was larger than the one evoked by serum from mice fed with SD (Fig. 3D, right panel). The amplitude of the current was reduced when the HFD-S was applied on DRG neurons from ASIC3 ko mice. These findings show that HFD-S activated depolarizing currents and induced hyperexcitability in DRGs cultured from WT mice, and that ASIC3 plays a significant role in this effect.

High-fat diet induced obesity is closely associated with dyslipidemia, which is characterized by an alteration in plasma lipid levels. Because dietary lipids can affect serum composition, we wanted to determine whether lipids in the serum of HFD-fed mice were responsible for the activation of ASIC3, leading to DRG hyperexcitability. Indeed, we have already identified lipids, in particular lysophosphatidylcholine (LPC16:0, LPC18:0 and LPC18:1) and arachidonic acid (AA), as endogenous activators of ASIC3 at physiological resting pH 7.4, *i.e.*, in the absence of extracellular acidification (Deval et al., 2008; Jacquot et al., 2021; Marra et al., 2016). LPC alone evokes depolarizing current in DRGs at physiological pH 7.4, increases skin nociceptive C-fiber firing, and induces both acute and chronic pain behaviors in rodents, depending on whether it is injected into the skin or joints (Jacquot et al., 2021; Marra et al., 2016; Pidoux et al., 2020). To test this hypothesis, we depleted lipids from HFD-serum (HFD-dl), and we perfused HFD-dl on WT-DRG neurons. The amplitude of the current evoked in DRG neurons during the application of HFD-dl serum was reduced compared to the full HFD-S (Fig. 3D, right panel). The amplitude of the current measured with HFD-dl was not different from that measured with SD-serum (Fig. 3D, right panel).

### HFD consumption led to an increase in LPC16:0, LPC18:0 and LPC18:1 concentration in serum from obese mice, which activated non-acidic ASIC3 currents

Experiments done in DRG cultures highlighted that lipids in HFD-S have a direct effect on DRG neurons. Accordingly, we then evaluated the impact of HFD on the lipid profile of serums from obese mice in order to identify lipid species that may act on ASIC3 channels leading to the increased neuronal excitability. Therefore, we studied the composition of serum from HFD- and SD-fed WT mice with lipidomic mass spectrometry analysis. We observed that the total amount of phosphatidylcholine and lysophosphatidylcholine species were elevated in the serum of HFD-fed mice compared to the serum of SD-fed mice (Supplementary Fig.3). In addition, LPC species were present in elevated concentration with major differences for LPC16:0, LPC18:0, LPC18:1, and LPC18:2 in HFD serum (Fig 4A). The concentration of LPC species were reduced significantly upon the process of delipidation of the HFD-S to levels that were similar to the SD-S (Fig. 4A). This observation was significant because lysophosphatidylcholine (LPC), together with arachidonic acid (AA), is the most efficient endogenous lipid activator of ASIC3 (Marra et al., 2016).

**Fig. 4.**
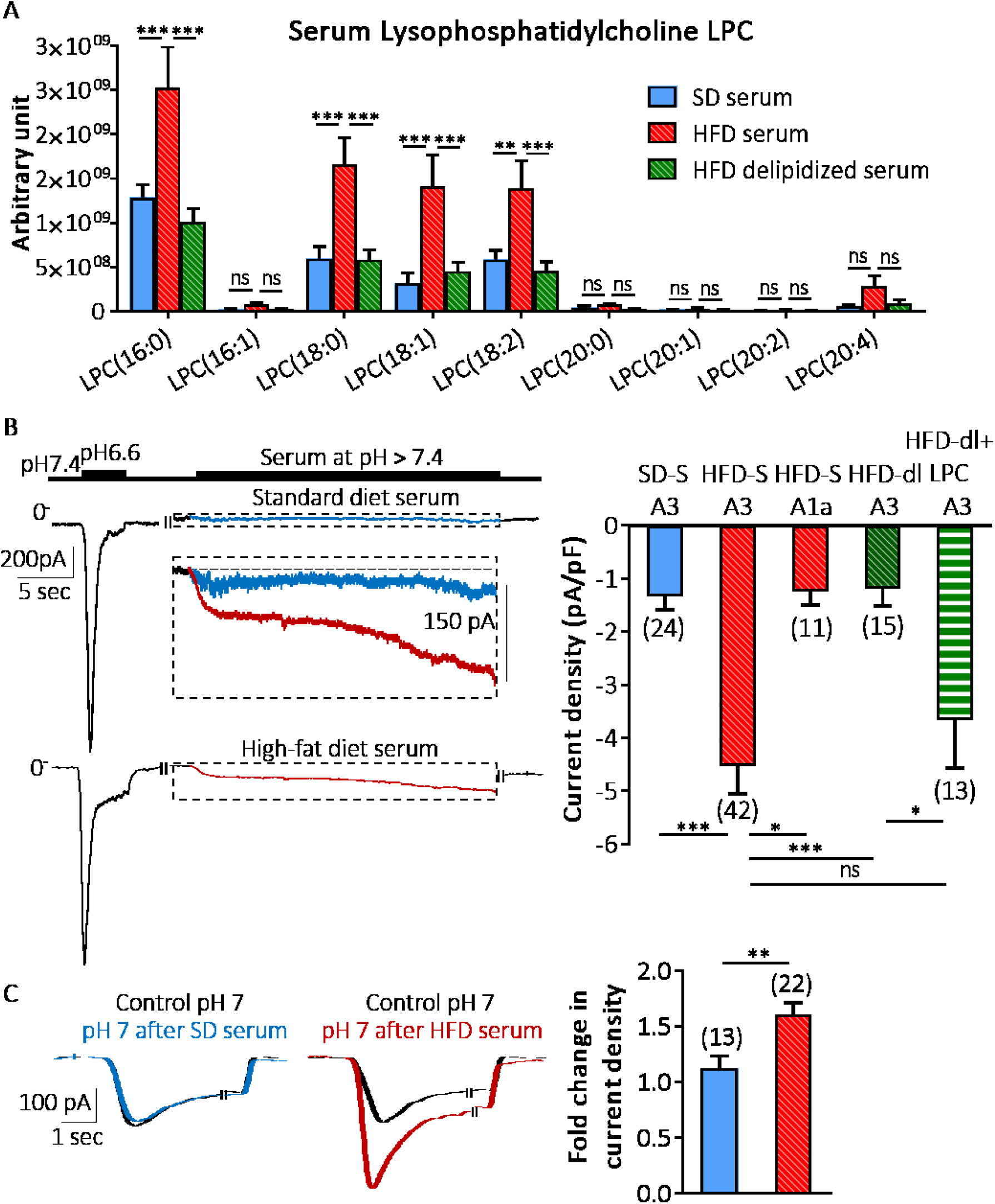
Lipid-rich diet increases LPC species in serum that activates and potentiates recombinant ASIC3 channels. **(A)** Bar diagram comparing the sum of quantitative responses (peak area signals) of different LPC species in serum collected from lean SD-fed and obese HFD-fed mice on diet for 12 weeks (F (16, 54) =4.3 p<0.0001, ns=not significant, **p<0.01, ***p<0.0001, 2-way ANOVA followed by Bonferroni’s for multiple comparison test, n=3 per group). **(B)** Left panel, representative recordings of the effect of SD-S (top trace, blue) and HFD-S (bottom trace, red) application on mouse ASIC3 channel expressed in HEK-293 cells, using whole-cell voltage-clamp configuration at −80 mV membrane potential. Inset shows a superimposed magnification of the original traces, as highlighted by dashed line squares. HFD-S directly activated ASIC3 channels compared to the effect of SD-S. Right panel shows the average effect of SD-S (current density −1.3 +/− 0.3 pA/pF), HFD-S (−4.5 +/− 0.5 pA/pF), HFD-dl (delipidized HFD-S; −1.2 +/− 0.3 pA/pF) and HFD-dl + LPC (HFD-dl supplemented with LPC16:0, LPC18:0 and LPC18:1; −3.7 +/− 0.9 pA/pF) application on ASIC3 current, and HFD-S on mouse ASIC1a current (current density −1.5 +/− 0.3 pA/pF, number of neurons recorded are indicated above each bar; current densities are measured 60 sec after serum application; p= 0.0001 for HFD-S vs SD-S, p= 0.02 for HFD-S on ASIC3 vs HFD-S on ASIC1a channels, p= 0.0002 HFD-S vs HFD-dl on ASIC3 channels, p > 0.99 for SD-S vs HFD-dl on ASIC3 and HFD-S on ASIC1a, p > 0.99 for HFD-S vs HFD-dl + LPC on ASIC3 channel; Kruskal-Wallis test followed by Dunn’s multiple comparisons test). **(C)** Effect of HFD-S and SD-S on ASIC3 current evoked at pH 7.0. Representative superimposed traces of ASIC3 currents activated at pH 7.0, showing control currents before serum application (black traces), the absence of effect of SD-S (blue trace), and the potentiation of ASIC3 currents at pH7.0 by HFD-S (red trace). The right panel presents the fold change in the current density pre vs post application of SD-S and HFD-S (n= 13-22 for SD-S and HFD-S respectively, p= 0.006; Unpaired t-test).

To test their direct effect on ASIC channels, serums were applied on HEK-293 cells expressing recombinant ASIC channels and currents were recorded with whole-cell patch-clamp technique in voltage-clamp configuration (Fig. 4B). The expression of ASIC channels was controlled on each cell with a 5 second application of an extracellular solution at pH 6.6 prior to the application of serum, which produced a transient inward current characteristic of the response of ASIC channels at acidic pH (Fig. 4B). In HEK-293 cells expressing mouse ASIC3 channel, the application of serum at neutral physiological pH7.4 activated a constitutive current similar to the current recorded in DRG cultures. Like in DRG cultures, currents were activated immediately after application of the serum, showing a gradual increase in amplitude, and were fully reversible upon withdrawal of the serum (Fig. 4B). The amplitude of the current measured with HFD-S was larger than with SD-S serum. ASIC1a, which is also expressed in DRGs (Deval & Lingueglia, 2015) was also tested. There was no effect of HFD-S on ASIC1a expressed in HEK-293 cells, and the evoked currents were not different from those measured with SD-S. ASIC1a is not activated by extracellular LPC (Jacquot et al., 2021; Marra et al., 2016; Pidoux et al., 2020), and these data are therefore consistent with a role of LPC present in HFD serum on ASIC3 activity in DRGs. We further investigated the contribution of lipids in HFD-S with HFD-dl serum applied on HEK-293 cells expressing ASIC3. The amplitude of the current activated with HFD-dl serum was reduced compared with HFD serum and it was not different from that activated with SD-S. We then complemented the lipid-depleted HFD serum with a cocktail of LPC species with concentrations corresponding to that measured in the full HFD-S (LPC16:0, LPC18:0, and LPC18:1, 20μM each). The application of supplemented HFD-dl serum on ASIC3 expressing HEK-293 cells rescued the current amplitude measured with HFD-S (Fig. 4B). The current activated by the application of HFD-S on non-transfected cells was not different from the one measured with SD-S on mASIC3 transfected cells or vehicle solution (Supplementary Fig. 4).

We then explored the effect of serum on acid-activated ASIC3 current at pH 7.0 (Fig. 4C). This pH is more acidic than the serum pH, so, to preserve the composition of the serum, HEK-293 cells were pre-incubated with HFD-S or SD-S for 1 min before re-application of pH 7.0 acid solution for 5 seconds. Acid-activated ASIC3 current was potentiated by pre-incubation with HFD-S (Fig. 4C). On the contrary, SD-S had no effect on ASIC3 current at pH 7.0.

Taken together, our data demonstrate that elevated lipid concentration in HFD serum, in particular LPC species, activates excitatory depolarizing ASIC3 current, leading to increased DRG neuron excitability.

### Genetic deletion and *in vivo* pharmacological inhibition of ASIC3 can prevent and correct HFD-induced thermal hyperalgesia

We next wanted to investigate the contribution of ASIC3 in heat pain hypersensitivity observed in HFD-fed obese mice. Two groups of ASIC3 ko mice were fed with either SD or HFD. ASIC3 ko mice fed with HFD developed obesity compared to ASIC3 ko mice fed with SD (Fig. 5A). Similar to our observation with HFD-fed WT mice, fasting blood glucose concentration of HFD-fed ASIC3 ko mice remained below 125 mg/dl during the first 12 weeks of diet and were not different from SD-fed mice (Supplementary Fig. 5A). Obese ASIC3 ko mice also showed progressive dysregulation of glucose homeostasis and increased insulin resistance. After 8 weeks of diet, the peak blood glucose concentration of HFD-fed ASIC3 ko mice was significantly higher than that measured in SD-fed mice (Supplementary Fig. 5B). At 12 weeks of HFD diet, blood glucose concentrations measured with GTT were not different between HFD-fed WT and ASIC3 ko mice (Supplementary Fig. 5C). This is consistent with a previous study that showed no difference in glucose regulation and insulin sensitivity between WT and ASIC3 inactivated conditions in young mice (Huang et al., 2008). Lipidomic analysis of serum from HFD-fed ASIC3 ko mice showed similar levels of LPC species compared to HFD-fed WT mice (Supplementary Fig. 6).

**Fig. 5.**
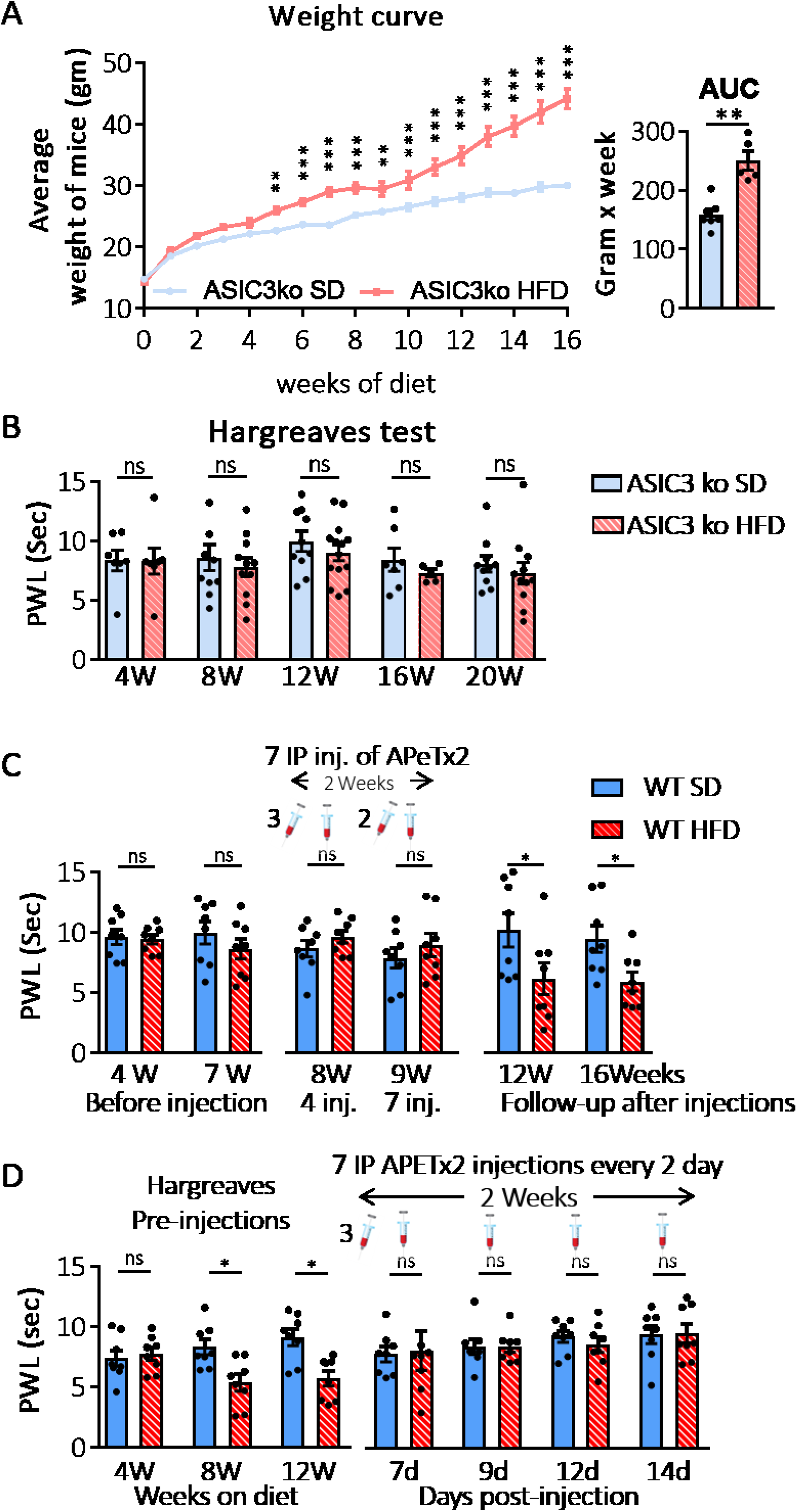
Genetic deletion and *in vivo* pharmacological inhibition of ASIC3 protect mice from lipid-rich diet induced thermal hyperalgesia. **(A)** Average weight of ASIC3 knockout mice fed with SD (pale blue) and HFD (pale red) (the interaction between diet and time, F (16, 176= 26.5, p<0.0001, n=7 mice per group, **p<0.01, ***p<0.001, 2-way ANOVA with repeated measures followed by Bonferroni’s post-hoc test). The right panel shows Area Under the Curve (p= 0.003 Mann-Whitney test). **(B)** Graph showing measures of heat perception with paw withdrawal latencies (PWL) from the Hargreaves test performed on ASIC3 knockout mice (ASIC3 ko) fed with SD (pale blue) and HFD (pale red). Weeks on diet (W) are indicated beneath the graph. The thermal perception was not significantly affected by diet in both groups (n=14-5, F (4, 60) = 0.08, p= 0.98, 2-way ANOVA with repeated measures followed by Bonferroni’s post-hoc test). **(C)** Effect of pharmacological inhibition of ASIC3 with APETx2 on the onset of heat hypersensitivity of HFD-fed WT mice. PWL measurements on SD-fed and HFD-fed WT mice that received 7 intraperitoneal injections (inj.) with the ASIC3 channel blocker APETx2 (0.23 mg/kg). A single injection was delivered every two days over 2 weeks, starting from the 7^th^ week of diet. At 4 and 7 weeks of diet, before starting the injections, both groups did not show any significance differences in the PWL. At 8 and 9 weeks of diet, during APETx2 treatment, the PWL was not statically significant (p= 0.9 for both timepoints). 3 and 7 weeks after the last APETx2 injection, at 12 and 16 weeks of diet, PWL was decreased in HFD-fed mice compared to SD-fed mice (n=8 per group, ns not significant, p=0.01 at 12 W and p=0.04 at 16 W of diet, 2 way ANOVA followed by Bonferroni’s multiple comparisons test). **(D)** Effect of pharmacological inhibition of ASIC3 with APETx2 on the heat hypersensitivity of HFD-fed WT mice, n=8 per group. PWL from the Hargreaves test performed on SD-fed and HFD-fed WT mice (p= 0.9 for SD vs HFD-fed mice at 4 W of diet, p= 0.04 and 0.01 for SD vs HFD-fed mice at 8 and 12 W of diet respectively). Then, mice received APETx2 intraperitoneal injections (0.23 mg/kg) using the same protocol used in (C), starting from 12 weeks of diet over two weeks (cartoon)(after 4 IP injections, 7 days of treatment and 13 weeks of diet, p=0.9 for HFD- vs SD-fed mice; p=0.9 after 9, 12 and 14 days of treatment, 2-way ANOVA followed by Bonferroni’s multiple comparisons test).

We then tested the heat sensitivity of obese HFD-fed ASIC3 ko mice. Interestingly, in contrast to the behavior of WT mice in the Hargreaves radian heat test, ASIC3 ko mice fed with HFD did not show heat hypersensitivity during 20 weeks of diet (Fig. 5B). Hind-paw withdrawal latencies of HFD-fed ASIC3 ko mice were not different from SD-fed ASIC3 ko mice (F (4, 60) = 0.08, p= 0.98, 2-way ANOVA with repeated measures followed by Bonferroni’s post-hoc test, n= 14-5). These data confirm the important role of ASIC3 channels in pain hypersensitivity to heat associated with HFD-induced obesity in mice.

We then investigated the effect of pharmacological inhibition of ASIC3 channels *in vivo*. Two groups of 8 WT mice at the age of 4 weeks were fed with SD and HFD for 7 weeks. Mice did not show heat hypersensitivity when tested with Hargreaves test at 4 and 7 weeks of diet (Fig. 5C). Then mice were IP injected every 2 days for 2 weeks with the ASIC3 inhibitor APETx2 (0.23 μg/g), starting after the Hargreaves test performed at week 7 of diet. These groups of mice were then tested for heat hypersensitivity on week 8 and 9 of diet. Heat sensitivity was not changed in the group of SD-fed mice (Fig. 5C). In contrast to the experiments performed in parallel with groups of HFD-fed WT mice that showed heat hypersensitivity after 8 weeks of diet (Fig. 2B), HFD-fed mice injected with ASIC3 inhibitor did not show heat hypersensitivity in the Hargreaves test at 8 or 9 weeks of diet (9.6 +/− 1.27 and 8.9 +/− 1.3 sec at 8 and 9 weeks, respectively, Fig. 5C). No additional injection was done on these mice and mice were kept on HFD or SD for 7 additional weeks. Heat hyperalgesia was then tested with Hargreaves test 3 and 7 weeks after the ultimate injection of ASIC3 inhibitor (*i.e.*, at 12 and 16 weeks of diet). The group of mice fed with SD did not show any variation of heat sensitivity over this period. The group of mice fed with HFD showed lower latencies indicating clear heat hypersensitivity compared to SD-fed mice (Fig. 5C). In another set of experiments, 2 groups of 4 weeks-old WT mice were fed with SD or HFD for 12 weeks. As previously observed (Fig. 2B), HFD-fed mice showed heat hypersensitivity in the Hargreaves test starting after 8 weeks of diet, while SD-fed mice did not show any change of their thermal sensitivity (Fig. 5D). Both groups of mice were then injected with ASIC3 inhibitor (APETx2, 0.23 μg/g), every two days over 2 weeks. This treatment did not change heat sensitivity of SD-fed mice, but it reversed heat hyperalgesia of HFD-fed mice and restored their perception to the same level as SD-fed mice (at 7 days’ post-injection PWL was 7.8 +/− 0.7 and 8.07 +/− 0.6 sec for SD and HFD mice respectively, p > 0.9, 2-way ANOVA with repeated measures followed by Bonferroni’s post-hoc test; Fig. 5D). All these data showed that pharmacological inhibition of ASIC3 can prevent and correct thermal hyperalgesia associated with the consumption of HFD, and support a role for ASIC3 in heat hyperalgesia associated with HFD.

## Discussion

The present study shows that high-fat diet-induced obesity in wild-type mice is associated with thermal heat hypersensitivity, which is correlated with an increased response to warming of heat-sensitive C-fibers in the skin and increased DRG neurons excitability. We show that serum from HFD-fed obese mice contains high concentrations of lipids, which depolarize small and medium diameter peripheral sensory DRG neurons, supposedly nociceptive neurons, and trigger action potential firing. This is the first demonstration that dietary lipids may affect the composition of serum to induce peripheral nociceptive DRG neurons hyperexcitability and pain through activation of an excitatory ion channel (ASIC3). The dyslipidemia and obesity associated with HFD consumption induce metabolic stress leading to the induction of several inflammatory signaling pathways within the metabolic tissue and especially the adipose tissue (Gregor & Hotamisligil, 2011). These signaling pathways elevate the release of several inflammatory cytokines, which decrease insulin signaling and affect glucose homeostasis (Hirosumi et al., 2002; Uysal et al., 1998). This well documented role of lipids in metabolic tissues can be extended to other organs and tissues that could be also affected. This is the case for the peripheral sensory nervous pathway that is dramatically affected by dysregulations in metabolic parameters that can cause peripheral diabetic neuropathies (Feldman et al., 2017). Nevertheless, the etiologies of pain associated with diabetic neuropathies are not well characterized. For example, several clinical studies showed that effective glycemic control can improve symptoms of diabetic neuropathy in type I but not in type II diabetes, therefore it is important to consider that different mechanisms and consequences are involved according to the models of diabetes (Callaghan et al., 2012).

Many studies on obesity and type 2 diabetes used genetic models, such as ob/ob or db/db mice respectively, obtained by interrupting leptin signaling pathway resulting in hyperphagia, obesity, and metabolic syndrome, including insulin resistance. However, genetic predispositions contribute to less than 10% of the physiological causes of obesity, which is far from representing the large population that suffers from obesity and its consequences (Le Thuc et al., 2017; Xu & Xue, 2016). The use of lipid rich diet in models of Diet Induced Obesity gives a strong advantage in being closer to the physiological causes of obesity in human, but some of the disadvantages of the DIO models for research on obesity and metabolic disorders are the severity of treatment needed to optimize the progression of obesity, which boosts its consequences and reduces inter-individual variability (O’Brien et al., 2014). The work presented here shows that consumption of lipid rich diet has harmful impacts on health, increasing the mice body weight and inducing obesity after a few weeks of diet. While HFD-fed mice progressively showed impaired glucose tolerance and insulin resistance over time, their fasting glucose did not exceed 150mg/dl (O’Brien et al., 2014; Obrosova et al., 2007) before 16 weeks of diet. In addition, insulin tolerance tests revealed that obese mice, regardless of genotype, showed significant insulin intolerance after 12 weeks of diet. Overall, these observations suggest that mice between 8 and 12 weeks of diet were in a pre-diabetic state. It was therefore important to investigate the neurophysiological consequences of a diabetogenic diet on the pre-diabetic state of obese mice between 8 weeks and 12 weeks of diet. The lipid-rich diet also induced dyslipidemia with increased concentration of several lipid species in the serum of obese mice. Contradictory results have been published in the literature about the plasma concentrations of LPC and AA associated with obesity (Barber et al., 2012; Fekete et al., 2015; Pickens et al., 2017; Pietiläinen et al., 2007). Indeed, it is important to consider that the composition of lipids in the serum strongly depends not only on the endogenous synthesis of fatty acids by adipose tissue but also on the intake of fatty acids present in the diet. We found elevated concentrations of lipids in the serum of HFD-fed obese mice, and interestingly, the main lipids species present in the high-fat diet (*i.e.* palmitic, stearic and oleic acids) were also found in conjugated forms in the serum phospholipids of obese mice. Therefore, dietary lipids in the HFD influence the composition of lipids in obese mice serum. It has recently been shown that enrichment of diet with ω-6 polyunsaturated fatty acids induces hyperexcitability of peripheral sensory neurons and pain in mice (Boyd et al., 2021). Although this study shows that high ω-6 intake can affect DRG neurons and pain, this cannot, however, explain the effect we observed because in the present study, the ω-6/ω-3 ratio in HFD diet was not higher than that in the SD (Supplementary table 1).

When applied onto recombinant ASIC3 channel, HFD serum activated the channel at non-acidic pH 7.4 with no delay, possibly indicating a direct activation. The HFD serum also potentiated the ASIC3 channel response to acidification to pH 7.0, without having an effect on ASIC1a channel. This effect on ASIC3 required the presence of lipid because delipidizing the serum strongly reduced its effect on ASIC3 channels. Weak acidification or hyperosmolarity could also activate ASIC3 currents (Deval et al., 2008), however, these do not seem to significantly account for the effect of HFD serum because both HFD and SD serum were measured at physiological neutral pH and limited hyperosmolarity. Although several species of lipids were significantly increased in the serum of mice fed with HFD, we focused on the role of LPC because it is one of the most concentrated lipid in serum of obese mice and we have shown previously that it activates and potentiates ASIC3 channel (Marra et al., 2016). The most concentrated LPCs in the serum (LPC16:0, LPC18:0 and LPC18:1) are also the most efficient lipids effective on ASIC3. Supplementing the delipidized serum with a cocktail of LPCs at a final concentration comparable to those found in HFD serum, partially restored the activity of the delipidized HFD serum on ASIC3 channels. These data strongly suggest that LPC found at high concentration in the serum of HFD mice can activate ASIC3 channel in DRG neurons. The inward current evoked by HFD serum in cultured DRG neurons was not observed in DRG neurons from ASIC3 knockout mice, indicating that the serum exerts its depolarizing effect primarily through the activation of ASIC3 channels. In addition, the excitatory effects of HFD serum on cultured DRG neurons (*i.e.*, activation of an inward depolarizing current and diminution of the rheobase) were reduced by pharmacological inhibition of ASIC3 with the peptide toxin APETx2 and in neurons from ASIC3 knockout mice.

We then confirmed *in vivo* the role of ASIC3 channels identified *in vitro* with DRG neurons. ASIC3 knockout mice fed with HFD were obese and showed deregulation in glucose homeostasis similarly to WT mice fed with the same diet, but they were protected from thermal heat-hypersensitivity. Lipidomic analysis of serum from obese ASIC3 ko mice showed similar levels of LPC species, compared to the serum from obese WT mice. This highlights that the protective effect, against thermal hypersensitivity associated with HFD consumption, observed in ASIC3 ko is therefore due to the loss of ASIC3 channels the mice and not to an alteration in the lipid composition of the serum of ASIC3 ko mice. Pharmacological studies with APETx2 toxin then confirmed the role of the ASIC3 channels in heat hypersensitivity. Interestingly, pharmacological inhibition of ASIC3 channels in wild-type mice with the peptide toxin APETx2 delayed the onset of the thermal heat hypersensitivity and reversed the hypersensitivity when it was established. This suggests that ASIC3 is “tonically” involved in thermal hyperalgesia in HFD-fed mice and that its inhibition is able to reverse the pain-related hypersensitivity of HFD-fed mice.

HFD-fed obese mice did not develop a mechanical hypersensitivity often associated with diabetic neuropathy (Bierhaus et al., 2012; Tsantoulas et al., 2017). This could be due to the specificity in the diet composition (*e.g.*, the quantity and quality of lipids in the diet such as saturated vs unsaturated lipids, and ω-6/ω-3 ratio), the duration of the regime, and the age of the mice at the beginning of the feeding period, as we fed young mice at the age of 4 weeks old, and the prediabetic state of HFD-fed mice. Mechanical hypersensitivity is thought to be mediated through distal sensory neuropathy associated with myelinated Aβ fibers while we show here that the thermal hypersensitivity is associated with the sensitization of unmyelinated nociceptive C-fibers (Baron & Maier, 1995; Khan et al., 2002). Although ASIC3 is co-expressed with the heat sensor TRPV1 channel in small diameter DRG neurons, ASIC3 channels is also expressed in mechanosensitive fibers (Emery & Ernfors, 2018; Usoskin et al., 2015), and there are evidence for the role of ASIC3 in mechanoperception in tissues like the skin and muscle (Lin et al., 2016; Price et al., 2001). In ASIC3 ko mice, however, no deficit of mechanoperception was observed in C-mechanonociceptive fibers, with von Frey filaments and skin stretch, while, on the contrary, the sensitivity of myelinated mechanonociceptive Aδ fibers (AM fibers) was reduced (Price et al., 2001). One could hypothesize that channels expressed in the myelinate A fibers may be more protected by the myelin sheet from the increase of lipid components in serum after HFD consumption, especially LPC that can integrate into cell membranes (Plemel et al., 2018), therefore limiting the access of these lipid components from the serum to the channels (Feldman et al., 2017).

In conclusion, our experiments shed light on the severe impact of metabolic dysregulation induced by the consumption of fat-rich diets on the peripheral nervous system and pain, and on the role of ASIC3 channels expressed in nociceptive sensory neurons in these chronic pain conditions. They also highlight, together with our recent observation on rheumatic diseases (Jacquot et al., 2021) as well as publication on fibromyalgia (Hung et al., 2020) or mechanical pain (Sadler et al., 2021), the growing and critical role of LPC in chronic pain conditions, including in humans, through pain-related ion channels such as ASIC3. These data could therefore open very interesting clinical perspectives although more work would be needed to further explore the role of ASIC3 channels in pain in humans, and in disorders associated with metabolic syndromes in obese patients.

## References

Afshin, A., Forouzanfar, M. H., Reitsma, M. B., Sur, P., Estep, K., Lee, A., Marczak, L., Mokdad, A. H., Moradi-Lakeh, M., Naghavi, M., Salama, J. S., Vos, T., Abate, K. H., Abbafati, C., Ahmed, M. B., Al-Aly, Z., Alkerwi, A., Al-Raddadi, R., Amare, A. T., … Murray, C. J. L. (2017). Health Effects of Overweight and Obesity in 195 Countries over 25 Years. The New England Journal of Medicine, 377(1), 13–27. https://doi.org/10.1056/NEJMoa1614362

Ayala, J. E., Samuel, V. T., Morton, G. J., Obici, S., Croniger, C. M., Shulman, G. I., Wasserman, D. H., McGuinness, O. P., & NIH Mouse Metabolic Phenotyping Center Consortium. (2010). Standard operating procedures for describing and performing metabolic tests of glucose homeostasis in mice. Disease Models & Mechanisms, 3(9-10), 525–534. https://doi.org/10.1242/dmm.006239

Barber, M. N., Risis, S., Yang, C., Meikle, P. J., Staples, M., Febbraio, M. A., & Bruce, C. R. (2012). Plasma lysophosphatidylcholine levels are reduced in obesity and type 2 diabetes. PloS One, 7(7), e41456. https://doi.org/10.1371/journal.pone.0041456

Bardoni, R., Tawfik, V. L., Wang, D., François, A., Solorzano, C., Shuster, S. A., Choudhury, P., Betelli, C., Cassidy, C., Smith, K., de Nooij, J. C., Mennicken, F., O’Donnell, D., Kieffer, B. L., Woodbury, C. J., Basbaum, A. I., MacDermott, A. B., & Scherrer, G. (2014). Delta Opioid Receptors Presynaptically Regulate Cutaneous Mechanosensory Neuron Input to the Spinal Cord Dorsal Horn. Neuron, 81(6), 1312–1327. https://doi.org/10.1016/j.neuron.2014.01.044

Baron, R., & Maier, C. (1995). Painful neuropathy: C-nociceptor activity may not be necessary to maintain central mechanisms accounting for dynamic mechanical allodynia. The Clinical Journal of Pain, 11(1), 63–69.

Bierhaus, A., Fleming, T., Stoyanov, S., Leffler, A., Babes, A., Neacsu, C., Sauer, S. K., Eberhardt, M., Schnölzer, M., Lasitschka, F., Lasischka, F., Neuhuber, W. L., Kichko, T. I., Konrade, I., Elvert, R., Mier, W., Pirags, V., Lukic, I. K., Morcos, M., … Nawroth, P. P. (2012). Methylglyoxal modification of Nav1.8 facilitates nociceptive neuron firing and causes hyperalgesia in diabetic neuropathy. Nature Medicine, 18(6), 926–933. https://doi.org/10.1038/nm.2750

Bligh, E. G., & Dyer, W. J. (1959). A rapid method of total lipid extraction and purification. Canadian Journal of Biochemistry and Physiology, 37(8), 911–917. https://doi.org/10.1139/o59-099

Boyd, J. T., LoCoco, P. M., Furr, A. R., Bendele, M. R., Tram, M., Li, Q., Chang, F.-M., Colley, M. E., Samenuk, G. M., Arris, D. A., Locke, E. E., Bach, S. B. H., Tobon, A., Ruparel, S. B., & Hargreaves, K. M. (2021). Elevated dietary ω-6 polyunsaturated fatty acids induce reversible peripheral nerve dysfunction that exacerbates comorbid pain conditions. Nature Metabolism, 3(6), 762–773. https://doi.org/10.1038/s42255-021-00410-x

Callaghan, B. C., Hur, J., & Feldman, E. L. (2012). Diabetic Neuropathy: One disease or two? Current Opinion in Neurology, 25(5), 536–541. https://doi.org/10.1097/WCO.0b013e328357a797

Deuis, J. R., Dvorakova, L. S., & Vetter, I. (2017). Methods Used to Evaluate Pain Behaviors in Rodents. Frontiers in Molecular Neuroscience, 10, 284. https://doi.org/10.3389/fnmol.2017.00284

Deval, E., Baron, A., Lingueglia, E., Mazarguil, H., Zajac, J.-M., & Lazdunski, M. (2003). Effects of neuropeptide SF and related peptides on acid sensing ion channel 3 and sensory neuron excitability. Neuropharmacology, 44(5), 662–671. https://doi.org/10.1016/S0028-3908(03)00047-9

Deval, E., & Lingueglia, E. (2015). Acid-Sensing Ion Channels and nociception in the peripheral and central nervous systems. Neuropharmacology, 94, 49–57. https://doi.org/10.1016/j.neuropharm.2015.02.009

Deval, E., Noël, J., Gasull, X., Delaunay, A., Alloui, A., Friend, V., Eschalier, A., Lazdunski, M., & Lingueglia, E. (2011). Acid-sensing ion channels in postoperative pain. The Journal of Neuroscience: The Official Journal of the Society for Neuroscience, 31(16), 6059–6066. https://doi.org/10.1523/JNEUROSCI.5266-10.2011

Deval, E., Noël, J., Lay, N., Alloui, A., Diochot, S., Friend, V., Jodar, M., Lazdunski, M., & Lingueglia, E. (2008). ASIC3, a sensor of acidic and primary inflammatory pain. The EMBO Journal, 27(22), 3047–3055. https://doi.org/10.1038/emboj.2008.213

Diochot, S., Baron, A., Rash, L. D., Deval, E., Escoubas, P., Scarzello, S., Salinas, M., & Lazdunski, M. (2004). A new sea anemone peptide, APETx2, inhibits ASIC3, a major acid-sensitive channel in sensory neurons. The EMBO Journal, 23(7), 1516–1525. https://doi.org/10.1038/sj.emboj.7600177

Emery, E. C., & Ernfors, P. (2018). Dorsal Root Ganglion Neuron Types and Their Functional Specialization. The Oxford Handbook of the Neurobiology of Pain. https://doi.org/10.1093/oxfordhb/9780190860509.013.4

Fekete, K., Györei, E., Lohner, S., Verduci, E., Agostoni, C., & Decsi, T. (2015). Long-chain polyunsaturated fatty acid status in obesity: A systematic review and meta-analysis. Obesity Reviews, 16(6), 488–497. https://doi.org/10.1111/obr.12280

Feldman, E. L., Nave, K.-A., Jensen, T. S., & Bennett, D. L. H. (2017). New Horizons in Diabetic Neuropathy: Mechanisms, Bioenergetics, and Pain. Neuron, 93(6), 1296–1313. https://doi.org/10.1016/j.neuron.2017.02.005

Graessler, J., Schwudke, D., Schwarz, P. E. H., Herzog, R., Shevchenko, A., & Bornstein, S. R. (2009). Top-down lipidomics reveals ether lipid deficiency in blood plasma of hypertensive patients. PloS One, 4(7), e6261. https://doi.org/10.1371/journal.pone.0006261

Gregor, M. F., & Hotamisligil, G. S. (2011). Inflammatory Mechanisms in Obesity. Annual Review of Immunology, 29(1), 415–445. https://doi.org/10.1146/annurev-immunol-031210-101322

Hainsworth, K. R., Davies, W. H., Khan, K. A., & Weisman, S. J. (2009). Co-occurring Chronic Pain and Obesity in Children and Adolescents: The Impact on Health-related Quality of Life. The Clinical Journal of Pain, 25(8), 715–721. https://doi.org/10.1097/AJP.0b013e3181a3b689

Hirosumi, J., Tuncman, G., Chang, L., Görgün, C. Z., Uysal, K. T., Maeda, K., Karin, M., & Hotamisligil, G. S. (2002). A central role for JNK in obesity and insulin resistance. Nature, 420(6913), 333–336. https://doi.org/10.1038/nature01137

Hitt, H. C., McMillen, R. C., Thornton-Neaves, T., Koch, K., & Cosby, A. G. (2007). Comorbidity of obesity and pain in a general population: Results from the Southern Pain Prevalence Study. The Journal of Pain, 8(5), 430–436. https://doi.org/10.1016/j.jpain.2006.12.003

Hsu, W.-H., Lee, C.-H., Chao, Y.-M., Kuo, C.-H., Ku, W.-C., Chen, C.-C., & Lin, Y.-L. (2019). ASIC3-dependent metabolomics profiling of serum and urine in a mouse model of fibromyalgia. Scientific Reports, 9(1), 12123. https://doi.org/10.1038/s41598-019-48315-w

Huang, S.-J., Yang, W.-S., Lin, Y.-W., Wang, H.-C., & Chen, C.-C. (2008). Increase of insulin sensitivity and reversal of age-dependent glucose intolerance with inhibition of ASIC3. Biochemical and Biophysical Research Communications, 371(4), 729–734. https://doi.org/10.1016/j.bbrc.2008.04.147

Hung, C.-H., Lee, C.-H., Tsai, M.-H., Chen, C.-H., Lin, H.-F., Hsu, C.-Y., Lai, C.-L., & Chen, C.-C. (2020). Activation of acid-sensing ion channel 3 by lysophosphatidylcholine 16:0 mediates psychological stress-induced fibromyalgia-like pain. Annals of the Rheumatic Diseases, 79(12), 1644–1656. https://doi.org/10.1136/annrheumdis-2020-218329

Jacquot, F., Khoury, S., Labrum, B., Delanoe, K., Pidoux, L., Barbier, J., Delay, L., Bayle, A., Aissouni, Y., Barriere, D. A., Kultima, K., Freyhult, E., Hugo, A., Kosek, E., Ahmed, A. S., Jurczak, A., Lingueglia, E., Svensson, C. I., Breuil, V., … Deval, E. (2021). Lysophophatidyl-choline 16:0 mediates persistent joint pain through Acid-Sensing Ion Channel 3: Preclinical and clinical evidences. BioRxiv, 2021.03.29.437487. https://doi.org/10.1101/2021.03.29.437487

Karczewski, J., Spencer, R. H., Garsky, V. M., Liang, A., Leitl, M. D., Cato, M. J., Cook, S. P., Kane, S., & Urban, M. O. (2010). Reversal of acid-induced and inflammatory pain by the selective ASIC3 inhibitor, APETx2. British Journal of Pharmacology, 161(4), 950–960. https://doi.org/10.1111/j.1476-5381.2010.00918.x

Khan, G. M., Chen, S.-R., & Pan, H.-L. (2002). Role of primary afferent nerves in allodynia caused by diabetic neuropathy in rats. Neuroscience, 114(2), 291–299. https://doi.org/10.1016/S0306-4522(02)00372-X

Le Thuc, O., Stobbe, K., Cansell, C., Nahon, J.-L., Blondeau, N., & Rovère, C. (2017). Hypothalamic Inflammation and Energy Balance Disruptions: Spotlight on Chemokines. Frontiers in Endocrinology, 8, 197. https://doi.org/10.3389/fendo.2017.00197

Lin, S.-H., Cheng, Y.-R., Banks, R. W., Min, M.-Y., Bewick, G. S., & Chen, C.-C. (2016). Evidence for the involvement of ASIC3 in sensory mechanotransduction in proprioceptors. Nature Communications, 7(1), 11460. https://doi.org/10.1038/ncomms11460

Marcus, D. A. (2004). Obesity and the Impact of Chronic Pain. The Clinical Journal of Pain, 20(3), 186–191.

Marra, S., Ferru-Clément, R., Breuil, V., Delaunay, A., Christin, M., Friend, V., Sebille, S., Cognard, C., Ferreira, T., Roux, C., Euller-Ziegler, L., Noel, J., Lingueglia, E., & Deval, E. (2016). Non-acidic activation of pain-related Acid-Sensing Ion Channel 3 by lipids. The EMBO Journal, 35(4), 414–428. https://doi.org/10.15252/embj.201592335

Mills, S. E. E., Nicolson, K. P., & Smith, B. H. (2019). Chronic pain: A review of its epidemiology and associated factors in population-based studies. British Journal of Anaesthesia, 123(2), e273–e283. https://doi.org/10.1016/j.bja.2019.03.023

Noël, J., Zimmermann, K., Busserolles, J., Deval, E., Alloui, A., Diochot, S., Guy, N., Borsotto, M., Reeh, P., Eschalier, A., & Lazdunski, M. (2009). The mechano-activated K+ channels TRAAK and TREK-1 control both warm and cold perception. The EMBO Journal, 28(9), 1308–1318. https://doi.org/10.1038/emboj.2009.57

O’Brien, P. D., Sakowski, S. A., & Feldman, E. L. (2014). Mouse models of diabetic neuropathy. ILAR Journal / National Research Council, Institute of Laboratory Animal Resources, 54(3), 259–272. https://doi.org/10.1093/ilar/ilt052

Obrosova, I. G., Ilnytska, O., Lyzogubov, V. V., Pavlov, I. A., Mashtalir, N., Nadler, J. L., & Drel, V. R. (2007). High-Fat Diet–Induced Neuropathy of Pre-Diabetes and Obesity: Effects of “Healthy” Diet and Aldose Reductase Inhibition. Diabetes, 56(10), 2598–2608. https://doi.org/10.2337/db06-1176

Okifuji, A., & Hare, B. D. (2015). The association between chronic pain and obesity. Journal of Pain Research, 8, 399–408. https://doi.org/10.2147/JPR.S55598

Peigneur, S., Béress, L., Möller, C., Marí, F., Forssmann, W.-G., & Tytgat, J. (2012). A natural point mutation changes both target selectivity and mechanism of action of sea anemone toxins. FASEB Journal: Official Publication of the Federation of American Societies for Experimental Biology, 26(12), 5141–5151. https://doi.org/10.1096/fj.12-218479

Pereira, V., Busserolles, J., Christin, M., Devilliers, M., Poupon, L., Legha, W., Alloui, A., Aissouni, Y., Bourinet, E., Lesage, F., Eschalier, A., Lazdunski, M., & Noël, J. (2014). Role of the TREK2 potassium channel in cold and warm thermosensation and in pain perception. Pain, 155(12), 2534–2544. https://doi.org/10.1016/j.pain.2014.09.013

Pickens, C. A., Vazquez, A. I., Jones, A. D., & Fenton, J. I. (2017). Obesity, adipokines, and C-peptide are associated with distinct plasma phospholipid profiles in adult males, an untargeted lipidomic approach. Scientific Reports, 7, 6335. https://doi.org/10.1038/s41598-017-05785-0

Pidoux, L., Delanoe, K., Lingueglia, E., & Deval, E. (2020). ASIC3-dependent spinal cord nociceptive signaling in cutaneous pain induced by lysophosphatidyl-choline. BioRxiv, 2020.12.28.424561. https://doi.org/10.1101/2020.12.28.424561

Pietiläinen, K. H., Sysi-Aho, M., Rissanen, A., Seppänen-Laakso, T., Yki-Järvinen, H., Kaprio, J., & Orešič, M. (2007). Acquired Obesity Is Associated with Changes in the Serum Lipidomic Profile Independent of Genetic Effects – A Monozygotic Twin Study. PLoS ONE, 2(2), e218. https://doi.org/10.1371/journal.pone.0000218

Plemel, J. R., Michaels, N. J., Weishaupt, N., Caprariello, A. V., Keough, M. B., Rogers, J. A., Yukseloglu, A., Lim, J., Patel, V. V., Rawji, K. S., Jensen, S. K., Teo, W., Heyne, B., Whitehead, S. N., Stys, P. K., & Yong, V. W. (2018). Mechanisms of lysophosphatidylcholine-induced demyelination: A primary lipid disrupting myelinopathy. Glia, 66(2), 327–347. https://doi.org/10.1002/glia.23245

Poirot, O., Berta, T., Decosterd, I., & Kellenberger, S. (2006). Distinct ASIC currents are expressed in rat putative nociceptors and are modulated by nerve injury: ASIC function in rat sensory neurones. The Journal of Physiology, 576(1), 215–234. https://doi.org/10.1113/jphysiol.2006.113035

Price, M. P., McIlwrath, S. L., Xie, J., Cheng, C., Qiao, J., Tarr, D. E., Sluka, K. A., Brennan, T. J., Lewin, G. R., & Welsh, M. J. (2001). The DRASIC Cation Channel Contributes to the Detection of Cutaneous Touch and Acid Stimuli in Mice. Neuron, 32(6), 1071–1083. https://doi.org/10.1016/S0896-6273(01)00547-5

Ray, L., Lipton, R. B., Zimmerman, M. E., Katz, M. J., & Derby, C. A. (2011). Mechanisms of association between obesity and chronic pain in the elderly. Pain, 152(1), 53–59. https://doi.org/10.1016/j.pain.2010.08.043

Renaud, J. F., Scanu, A. M., Kazazoglou, T., Lombet, A., Romey, G., & Lazdunski, M. (1982). Normal serum and lipoprotein-deficient serum give different expressions of excitability, corresponding to different stages of differentiation, in chicken cardiac cells in culture. Proceedings of the National Academy of Sciences, 79(24), 7768–7772. https://doi.org/10.1073/pnas.79.24.7768

Rodgers, H. M., Liban, S., & Wilson, L. M. (2014). Attenuated pain response of obese mice (B6.Cg-lep(ob)) is affected by aging and leptin but not sex. Physiology & Behavior, 123, 80–85. https://doi.org/10.1016/j.physbeh.2013.10.007

Sadler, K. E., Moehring, F., Shiers, S. I., Laskowski, L. J., Mikesell, A. R., Plautz, Z. R., Brezinski, A. N., Mecca, C. M., Dussor, G., Price, T. J., McCorvy, J. D., & Stucky, C. L. (2021). Transient receptor potential canonical 5 mediates inflammatory mechanical and spontaneous pain in mice. Science Translational Medicine, 13(595), eabd7702. https://doi.org/10.1126/scitranslmed.abd7702

Song, Z., Xie, W., Chen, S., Strong, J. A., Print, M. S., Wang, J. I., Shareef, A. F., Ulrich-Lai, Y. M., & Zhang, J.-M. (2017). High-fat diet increases pain behaviors in rats with or without obesity. Scientific Reports, 7(1), 10350. https://doi.org/10.1038/s41598-017-10458-z

Tsantoulas, C., Laínez, S., Wong, S., Mehta, I., Vilar, B., & McNaughton, P. A. (2017). Hyperpolarization-activated cyclic nucleotide-gated 2 (HCN2) ion channels drive pain in mouse models of diabetic neuropathy. Science Translational Medicine, 9(409), eaam6072. https://doi.org/10.1126/scitranslmed.aam6072

Usoskin, D., Furlan, A., Islam, S., Abdo, H., Lönnerberg, P., Lou, D., Hjerling-Leffler, J., Haeggström, J., Kharchenko, O., Kharchenko, P. V., Linnarsson, S., & Ernfors, P. (2015). Unbiased classification of sensory neuron types by large-scale single-cell RNA sequencing. Nature Neuroscience, 18(1), 145–153. https://doi.org/10.1038/nn.3881

Uysal, K. T., Wiesbrock, S. M., & Hotamisligil, G. S. (1998). Functional analysis of tumor necrosis factor (TNF) receptors in TNF-alpha-mediated insulin resistance in genetic obesity. Endocrinology, 139(12), 4832–4838. https://doi.org/10.1210/endo.139.12.6337

Vos, T., Abajobir, A. A., Abate, K. H., Abbafati, C., Abbas, K. M., Abd-Allah, F., Abdulkader, R. S., Abdulle, A. M., Abebo, T. A., Abera, S. F., Aboyans, V., Abu-Raddad, L. J., Ackerman, I. N., Adamu, A. A., Adetokunboh, O., Afarideh, M., Afshin, A., Agarwal, S. K., Aggarwal, R., … Murray, C. J. L. (2017). Global, regional, and national incidence, prevalence, and years lived with disability for 328 diseases and injuries for 195 countries, 1990–2016: A systematic analysis for the Global Burden of Disease Study 2016. The Lancet, 390(10100), 1211–1259. https://doi.org/10.1016/S0140-6736(17)32154-2

Walder, R. Y., Rasmussen, L. A., Rainier, J. D., Light, A. R., Wemmie, J. A., & Sluka, K. A. (2010). ASIC1 and ASIC3 Play Different Roles in the Development of Hyperalgesia After Inflammatory Muscle Injury. The Journal of Pain, 11(3), 210–218. https://doi.org/10.1016/j.jpain.2009.07.004

Waldmann, R., Bassilana, F., Weille, J. de, Champigny, G., Heurteaux, C., & Lazdunski, M. (1997). Molecular Cloning of a Non-inactivating Proton-gated Na+ Channel Specific for Sensory Neurons. Journal of Biological Chemistry, 272(34), 20975–20978. https://doi.org/10.1074/jbc.272.34.20975

Webb, R., Brammah, T., Lunt, M., Urwin, M., Allison, T., & Symmons, D. (2003). Prevalence and predictors of intense, chronic, and disabling neck and back pain in the UK general population. Spine, 28(11), 1195–1202. https://doi.org/10.1097/01.BRS.0000067430.49169.01

Wu, W.-L., Cheng, C.-F., Sun, W.-H., Wong, C.-W., & Chen, C.-C. (2012). Targeting ASIC3 for pain, anxiety, and insulin resistance. Pharmacology & Therapeutics, 134(2), 127–138. https://doi.org/10.1016/j.pharmthera.2011.12.009

Wultsch, T., Painsipp, E., Shahbazian, A., Mitrovic, M., Edelsbrunner, M., Waldmann, R., Lazdunski, M., & Holzer, P. (2008). Deletion of the acid-sensing ion channel ASIC3 prevents gastritis-induced acid hyperresponsiveness of the stomach – brainstem axis. Pain, 134(3), 245–253. https://doi.org/10.1016/j.pain.2007.04.025

Xu, S., & Xue, Y. (2016). Pediatric obesity: Causes, symptoms, prevention and treatment. Experimental and Therapeutic Medicine, 11(1), 15–20. https://doi.org/10.3892/etm.2015.2853

Yan, J., Wei, X., Bischoff, C., Edelmayer, R. M., & Dussor, G. (2013). PH-evoked dural afferent signaling is mediated by ASIC3 and is sensitized by mast cell mediators. Headache, 53(8), 1250–1261. https://doi.org/10.1111/head.12152

Yen, Y.-T., Tu, P.-H., Chen, C.-J., Lin, Y.-W., Hsieh, S.-T., & Chen, C.-C. (2009). Role of acid-sensing ion channel 3 in sub-acute-phase inflammation. Molecular Pain, 5, 1. https://doi.org/10.1186/1744-8069-5-1

Zimmermann. (1983). Ethical guidelines for investigations of experimental pain in conscious animals. PAIN, 16(2), 109. https://doi.org/10.1016/0304-3959(83)90201-4

Zimmermann, Hein, A., Hager, U., Kaczmarek, J. S., Turnquist, B. P., Clapham, D. E., & Reeh, P. W. (2009). Phenotyping sensory nerve endings in vitro in the mouse. Nature Protocols, 4(2), 174–196. https://doi.org/10.1038/nprot.2008.223

