## Supplementary Figures for "Lysophosphatidylcholine induces heat pain hypersensitivity in obese mice fed with a high-fat diet through activation of peripheral Acid-Sensing Ion Channel 3"

Supplementary Table 1: Relative ratios of the main fatty acids found in standard and high-fat diets.

Supplementary Figure 1: Effect of HFD on glucose homeostasis in WT mice.

**(A**) Glucose tolerance test (GTT) for WT mice at 4 weeks of diet (glycemia at 120 min post injection was 159.2 +/- 6.6 mg/dl and 145.4 +/- 4.6 mg/dl for HFD- and SD-fed mice respectively, p=0.07, 2-way ANOVA with repeated measures followed by Bonferroni's post hoc test, n=10 per group; ns not significant, ***p<0.001, 2-way ANOVA with repeated measures followed by Bonferroni's post hoc test for multiple comparisons). Right panel shows AUC of individual replicates (p= 0.006, unpaired t-test). **(B)** GTT for WT mice at 8 weeks of diet (at 120 min post injection, blood glycemia was measured as 206.5 +/- 8.2 mg/dl and 153.8 +/- 5.4 mg/dl for HFD- and SD-fed mice respectively, p = 0.002, 2-way ANOVA with repeated measures followed by Bonferroni's post hoc test, n=10 per group). Right panel shows AUC of individual replicates (p=0.01 unpaired t-test) **(C)** GTT for WT mice at 16 weeks of diet (at 120 min post injection blood glycemia was measured as 134.7 +/-6 mg/dl and 265 +/- 26.5 mg/dl for SD- and HFD-fed mice respectively, n=10 per group, p< 0.0001, 2-way ANOVA with repeated measures followed by Bonferroni's post hoc test for multiple comparisons). Right panel shows AUC of individual replicates (p = 0.002, unpaired t-test).

Supplementary Figure 2: Mechanical sensitivity of obese HFD-fed WT mice at 16 weeks of diet.
Mechanical sensitivity was assessed with von Frey filaments of increasing stiffness (n= 10 per group, p= 0.7, unpaired t-test).

Supplementary Table 2: Cell membrane capacitances and resting potentials of cultured DRG neurons used to test the effect of serum on the rheobase.

Number (n) and mean values of membrane capacitance and of resting potential of DRG neurons (Kruskal-Wallis test).

Supplementary Table 3: Threshold of action potential triggering of cultured DRG neurons used to test the effect of serum on the rheobase.

Mean values of the threshold of action potential triggering during control rheobase and the rheobase during the application of either SD or HFD serum.

Supplementary Table 4: Cell membrane capacitance and resting potential of DRG neurons used to test the effect of serum on the whole-cell currents.

Number (n) and mean values of membrane capacitance and resting membrane potential of DRG neurons, Kruskal-Wallis test.

Supplementary Figure 3: Evaluation of the relative phospholipid and lysophospholipid composition of SD-S, HFD-S and HFD-delipidized with lipidomic mass spectrometric analysis.

Quantities of different lipid species found in the serum collected from SD-fed and HFD-fed mice, presented in arbitrary unit. PC (Phosphatidylcholine), PE (Phosphatidylethanolamine), PI (Phosphatidylinositol), LPC (Lysophosphatidylcholine), LPE (Lysophosphatidylethanolamine), p-value obtained from one-way ANOVA followed by Bonferroni's post hoc test; ns= not significant, *P < 0.05, **P ≤ 0.01.

### Supplementary Figure 4: Effect of HFD-serum on non-transfected HEK-293 cells, and of vehicle of delipidized HFD-serum complemented with LPC on HEK-293 cells expressing mouse ASIC3.

Calculated membrane current densities (pA/pF) obtained with patch-clamp recordings in voltage-clamp configuration in HEK-293 cells upon application of HFD serum (HFD-S) on non-transfected HEK-293 cells (current density -0.4 +/- 0.18 pA/pF, n= 8, p= 0.94 vs SD-S, Kruskal-Wallis test followed by a Dunn's post hoc). White bar represents the application of delipidized HFD serum (HFD-dl) supplemented with 0.12% EtOH (solvent used to dilute the synthesized LPC) on HEK-293 cells transfected with mouse ASIC3 channels, (current density -0.35 +/- 0.33 pA/pF, n= 8, p > 0.99 vs SD-S, Kruskal-Wallis test followed by a Dunn's post hoc).

### Supplementary Figure 5: HFD induces glucose intolerance and resistance to insulin in WT and ASIC3 knockout mice.

**(A)** Fasting blood glucose concentration measured at 4, 8, and 12 weeks of HFD of WT (red) and ASIC3 ko (pale red) mice (n=6, F (2, 18) = 1.9, p=0.2, 2-way ANOVA with repeated measures followed by Bonferroni's post-hoc test). **(B)** GTT performed at 8 weeks of diet in ASIC3 ko mice fed with SD and HFD (n= 7-5, respectively, ***p<0.001, 2-way ANOVA with repeated measures followed by Bonferroni's post-hoc test). The right panel shows AUC (p= 0.03, Mann-Whitney test). **(C)** GTT performed at 12 weeks of diet WT-HFD (red) and ASIC3 ko-HFD (pale red) (n= 5-6 respectively; not significant, 2-way ANOVA with repeated measures followed by Bonferroni's post-hoc test). The right panel shows AUC (p= 0.2, Mann-Whitney test). **(D)** ITT performed at 12 weeks of diet with SD-fed and HFD-fed WT (blue and red) and SD- and HFD-fed ASIC3 ko mice (pale blue and pale red) (n= 9-5, respectively; 2-way ANOVA with repeated measures followed by Bonferroni's post-hoc test. For comparison between WT -SD -HFD significance is represented as *P < 0.05, **P ≤ 0.01, ***P ≤ 0.001, while for ASIC3 ko -SD -HFD $^{\#}$P < 0.05, $^{\#\#}$P ≤ 0.01, $^{\#\#\#}$P ≤ 0.001).

### Supplementary Figure 6: HFD consumption increase serum LPC concentration in WT as well as ASIC3 knockout mice.

Bar diagram comparing the sum of quantitative responses (peak area signals) of different LPC species in serum collected from obese HFD-fed WT and ASIC3 knockout mice. Serum from both mice groups showed elevated LPC 16:0, 18:0, 18:1 and 18:2 that were not significantly different between groups (F (8, 36) = 1.5, p=0.2, 2-way ANOVA followed by Bonferroni's for multiple comparison test, n=3 per group).

Supplementary Table 1

| | | SD | HFD |
|---|---|---|---|
| **Total fat (%)** | | 5.1 | 34.2 |
| **Saturated FA (%)** | | 18.5 | 68.9 |
| **Unsaturated FA (%)** | | 81.4 | 31.1 |
| **Poly-unsaturated FA (%)** | | 59.7 | 7.9 |
| **Myristic acid (%)** | C14:0 | 0.4 | 10.7 |
| **Pentadecanoïc acid (%)** | C15:0 | 0.1 | 1.8 |
| **Palmitic acid (%)** | C16:0 | 14.6 | 33 |
| **Palmitoleic acid (%)** | C16:1 | 0.5 | 1.5 |
| **Stearic acid (%)** | C18:0 | 2.7 | 10.2 |
| **Oleic acid (%)** | C18:1 | 17.9 | 20 |
| **Linoleic acid (LA) (%)** | C18:2 | 50.8 | 5.7 |
| **Linolenic acid (ALA) (%)** | C18:3 | 5.5 | 0.9 |
| **Gondoic acid (%)** | C20:1 | 1.1 | 0.2 |
| **Arachidonic acid (%)** | C20:4 | 0.2 | 0.1 |
| **ω-3** | | 8.1 | 1 |
| **ω-6** | | 51.4 | 5.8 |
| **ratio LA/ALA** | | 9.2 | 6.3 |
| **ratio ω-6/ω-3** | | 6.3 | 5.6 |
| **ratio saturated FA/ω-3** | | 2.3 | 66.2 |
| **ratio C18:1/C16:0** | | 1.4 | 0.6 |

Supplementary figures 1

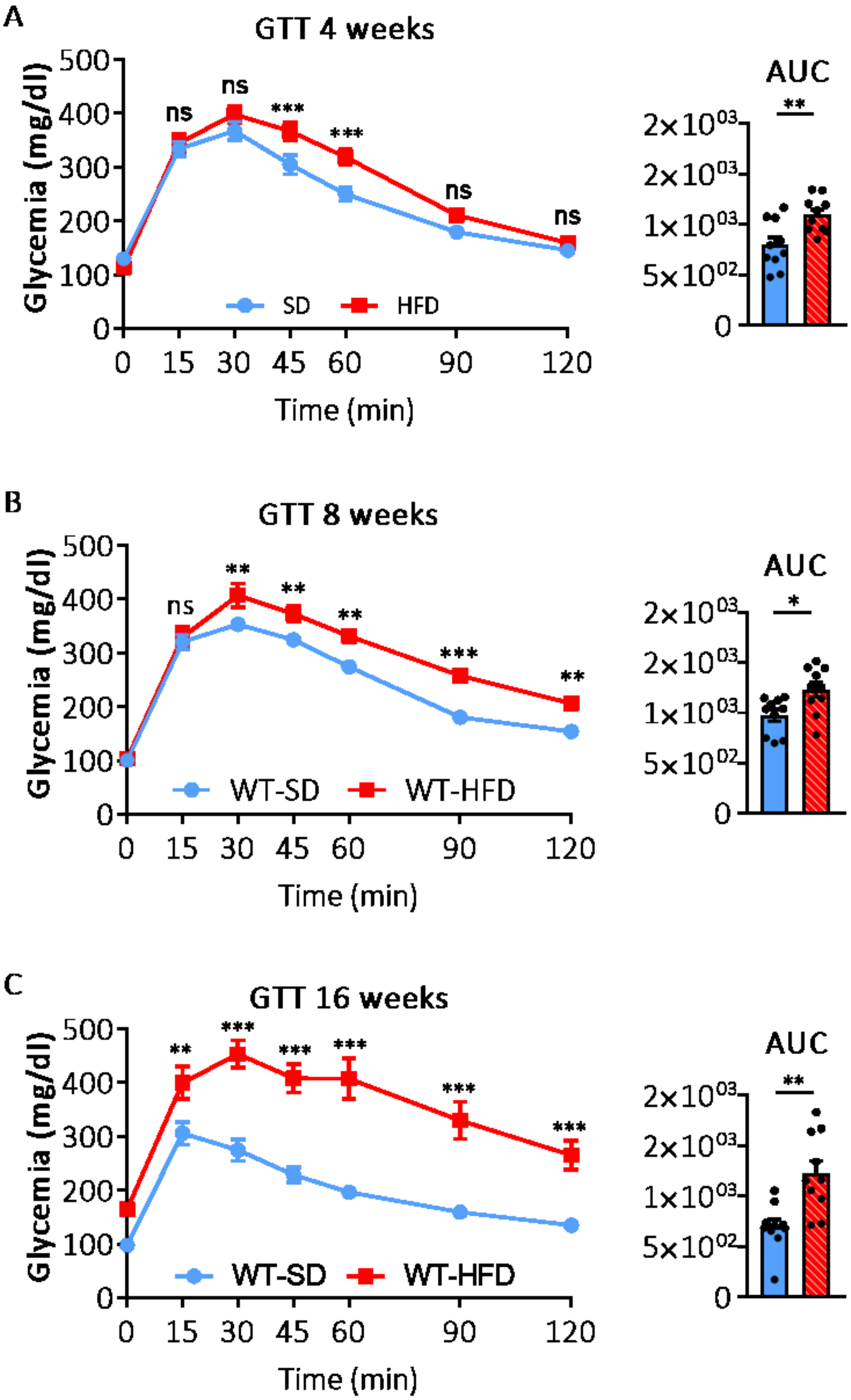

Supplementary figures 2

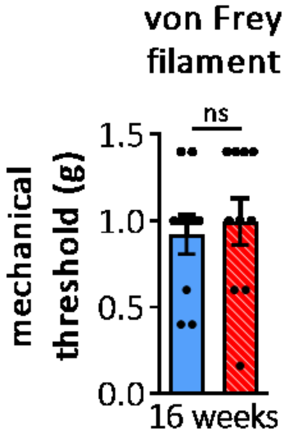

### Supplementary Table 2

| | SD serum on WT-DRG | HFD serum on WT-DRG | HFD serum on ASIC3 ko-DRG | Kruskal-Wallis test. |
|---|---|---|---|---|
| n | 8 | 11 | 15 | |
| Capacitance (pF) | 48.3±4.8 | 42.3±3.2 | 56.7±6.3 | F (2, 31) = 2.8 P=0.25 |
| Resting potential(mV) | -54.7±3.9 | -54.2±3.7 | -52.5±1.5 | F (2, 31) = 0.5 p= 0.76 |

### Supplementary Table 3

| | SD serum on WT-DRG | HFD serum on WT-DRG | HFD serum on ASIC3 ko-DRG |
|---|---|---|---|
| Threshold of AP at control rheobase(mV) | -10.5+/-2.3 | -6.6+/-3.6 | -9.9+/-1.6 |
| Threshold of AP during serum application (mV) | -8.8+/-1.5 | -11.53+/-3.9 | -8.9+/-1.6 |
| Threshold of AP at control rheobase vs during serum application Wilcoxon matched-pairs signed rank test | p= 0.2 | p= 0.1 | p=0.6 |

### Supplementary Table 4

| | SD serum on WT-DRG | HFD serum on WT-DRG | HFD serum on ASIC3-DRG | HFD-dl serum on WT-DRG | Kruskal-Wallis test. |
|---|---|---|---|---|---|
| n | 10 | 15 | 18 | 11 | |
| Capacitance | 35.2±3.2 | 33.2±3.7 | 33.1±2.6 | 34.5±4.5 | F (3, 50) = 0.87 p= 0.8 |
| Resting potential | -61.7±2.6 | -63.0±1.1 | -61.3±1.5 | -62.4±1.8 | F (3, 50) = 1.7 p= 0.63 |

Supplementary figures 3

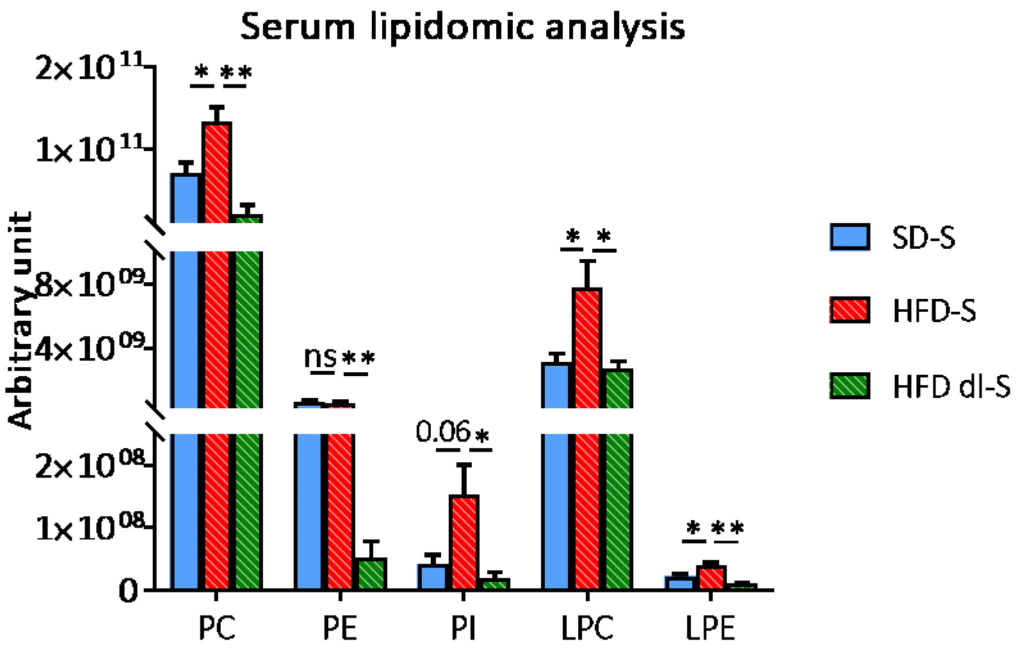

Supplementary figures 4

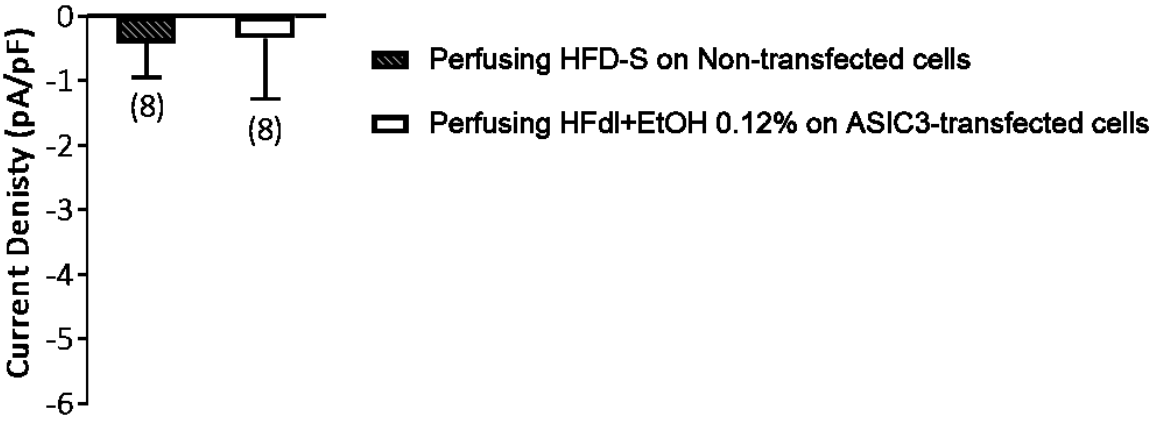

### Supplementary figures 5

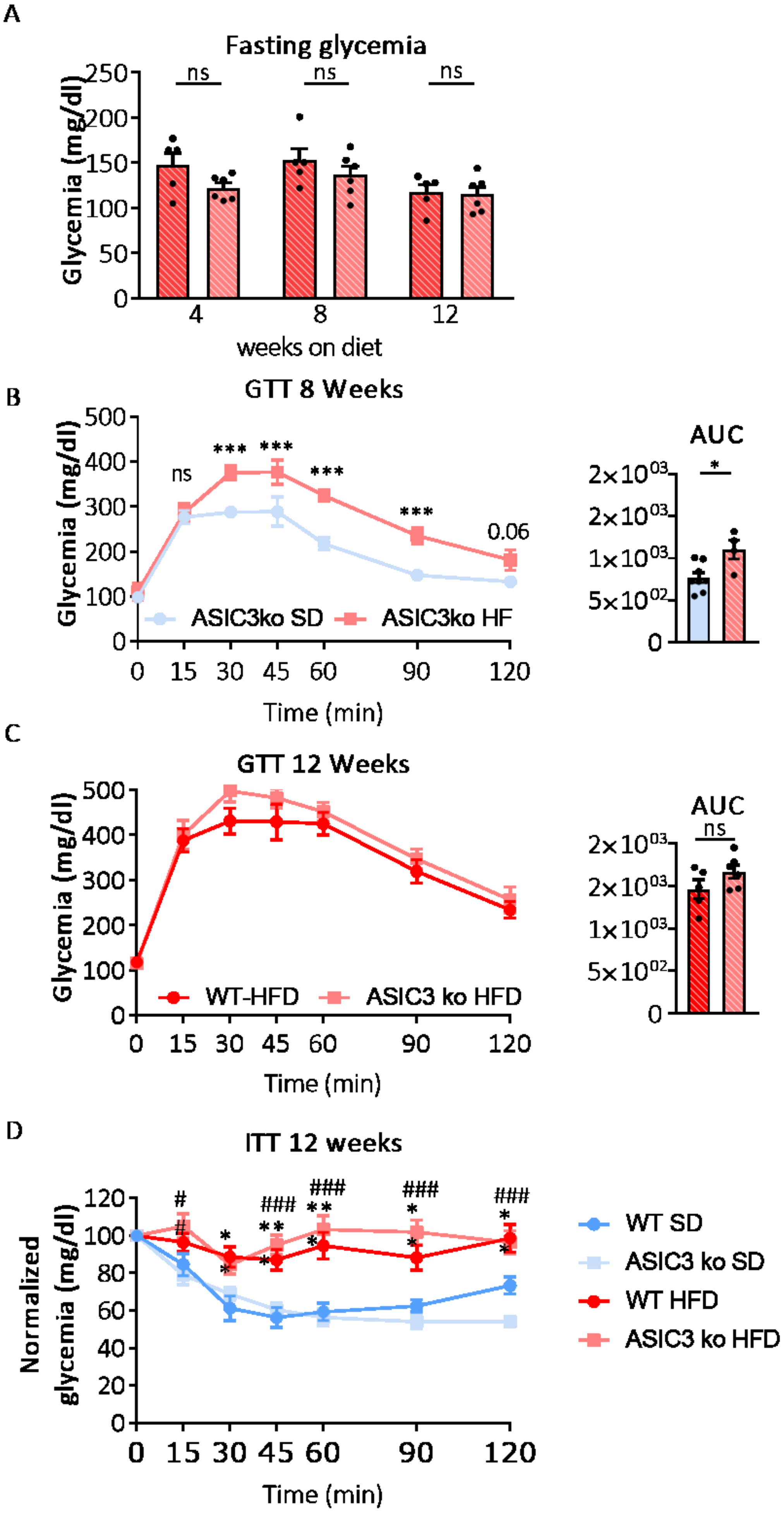

Supplementary figures 6

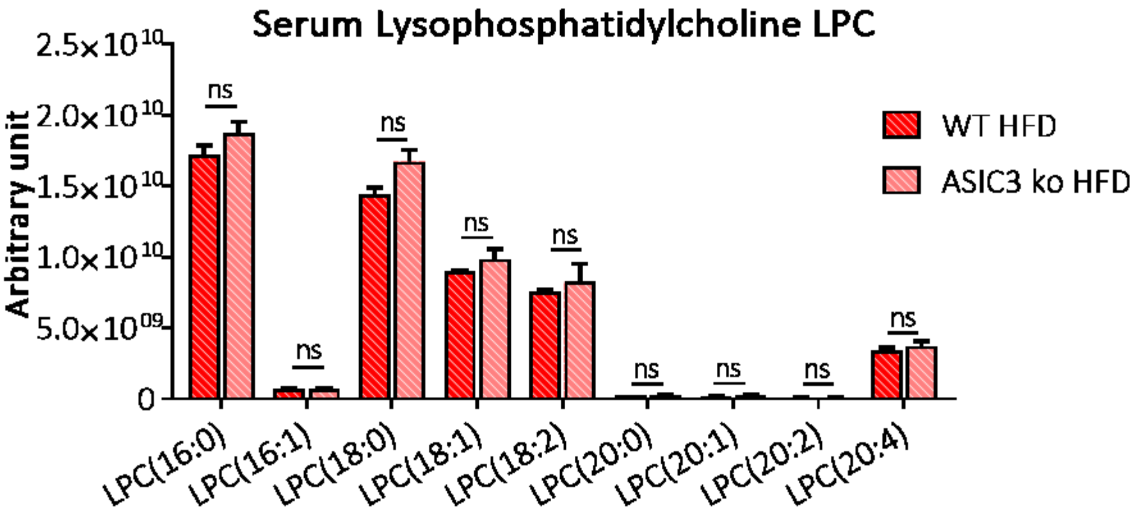
